# Identification and characterization of OmpT-like proteases in uropathogenic *Escherichia coli* clinical isolates

**DOI:** 10.1101/594564

**Authors:** Isabelle Desloges, James A. Taylor, Jean-Mathieu Leclerc, John R. Brannon, Andrea Portt, John D. Spencer, Ken Dewar, Gregory T Marczynski, Amee Manges, Samantha Gruenheid, Hervé Le Moual, Jenny-Lee Thomassin

**Affiliations:** Department of Microbiology and Immunology, McGill University, Montreal, QC, Canada; Division of Nephrology, Nationwide Children’s Hospital, Columbus, OH, USA; Microbiome and Disease Tolerance Centre, McGill University, Montreal, QC, Canada; Department of Human Genetics, McGill University, Montreal, QC, Canada; School of Population and Public Health, University of British Columbia, Vancouver, BC, Canada; Faculty of Dentistry, McGill University, Montreal, QC, Canada; Department of Structural Biology and Chemistry, Institut Pasteur, Paris, France; Department of Cellular and Molecular Medicine, University of Ottawa, Ottawa, ON, Canada; Dalla Lana School of Public Health, University of Toronto, Toronto, ON, Canada

**Keywords:** OmpT, OmpP, ArlC, antimicrobial peptides, RNase 7, LL-37, UPEC

## Abstract

Bacterial colonization of the urogenital tract is limited by innate defenses, including the production of antimicrobial peptides (AMPs). Uropathogenic *Escherichia coli* (UPEC) resist AMP-killing to cause a range of urinary tract infections (UTIs) including asymptomatic bacteriuria, cystitis, pyelonephritis, and sepsis. UPEC strains have high genomic diversity and encode numerous virulence factors that differentiate them from non-UTI causing strains, including *ompT*. As OmpT homologues cleave and inactivate AMPs, we hypothesized that high OmpT protease activity-levels contribute to UPEC colonization during symptomatic UTIs. Therefore, we measured OmpT activity in 58 UPEC clinical isolates. While heterogeneous OmpT activities were observed, OmpT activity was significantly greater in UPEC strains isolated from patients with symptomatic infections. Unexpectedly, UPEC strains exhibiting the greatest protease activities harboured an additional *ompT*-like gene called *arlC* (*ompTp*). The presence of two *OmpT*-like proteases in some UPEC isolates led us to compare the substrate specificities of OmpT-like proteases found in *E. coli*. While all three cleaved AMPs, cleavage efficiency varied on the basis of AMP size and secondary structure. Our findings suggest the presence ArlC and OmpT in the same UPEC isolate may confer a fitness advantage by expanding the range of target substrates.

## 1 Introduction

Urinary tract infections (UTIs) are among the most common cause of bacterial infections requiring antibiotic treatment (Flores-Mireles, Walker, Caparon, & Hultgren, 2015; Foxman, 2014; Hooton & Stamm, 1997). The majority of community acquired UTIs (70-95%) and recurrent UTIs are caused by uropathogenic *Escherichia coli* (UPEC) (Flores-Mireles et al., 2015; Nielubowicz & Mobley, 2010). The human gut acts as a reservoir for UPEC strains where they form part of the fecal flora (Kaper, Nataro, & Mobley, 2004; Moreno et al., 2006). Following colonization of the periurethral area, UPEC infect the urinary tract in an ascending manner, resulting in diseases ranging from asymptomatic bacteriuria (ABU), cystitis, pyelonephritis and sepsis (Hooton, 2012). UPEC strains have high genomic diversity and encode numerous virulence factors that differentiate them from non-UTI causing strains (Johnson, 1991; Lloyd, Rasko, & Mobley, 2007; Najafi, Hasanpour, Askary, Aziemzadeh, & Hashemi, 2018; Norinder, Koves, Yadav, Brauner, & Svanborg, 2012). These virulence factors contribute to disease progression allowing UPEC to colonize the uroepithelium, produce toxins, scavenge metabolites, and evade the host immune system (Schwab, Jobin, & Kurts, 2017; Terlizzi, Gribaudo, & Maffei, 2017).

Bacterial colonization is limited in the upper urogenital tract by several mechanisms including urine flow, chemical properties of urine, epithelial cell shedding, influx of immune cells including neutrophils upon bacterial stimulation, and secretion of soluble proteins and peptides by epithelial cells (Spencer, Schwaderer, Becknell, Watson, & Hains, 2014; Weichhart, Haidinger, Horl, & Saemann, 2008). Secreted proteins and antimicrobial peptides (AMPs) form part of the innate immune defenses of the urogenital tract and act through immunomodulation, indirect anti-colonization activity or direct bacterial killing (Kai-Larsen et al., 2010; Zasloff, 2007). AMPs are small (12-50 amino acids), cationic, amphipathic peptides that exert bactericidal action by interacting with anionic bacterial membranes to form pores resulting in bacterial lysis (Jenssen, Hamill, & Hancock, 2006). Two types of AMPs are detected in the urogenital tract: defensins that form small disulfide bond stabilized ß-sheets and the α-helical cathelicidin LL-37 (Chromek et al., 2006; Lehmann et al., 2002; Valore et al., 1998). In addition, the urogenital tract produces large structured antimicrobial proteins called ribonucleases (RNase) (Spencer et al., 2011; Spencer et al., 2013). Human α-defensin 5 (HD5), human ß-defensins (hBD) 1 and 2, LL-37 and RNase 7 are thought to prevent bacterial colonization as they are constitutively expressed in the urinary tract (Kjolvmark, Akesson, & Pahlman, 2017; Spencer et al., 2012). During UTIs HD5, hBD2, LL-37 and RNase 7 production increases, suggesting an active role in bacterial clearance (Chromek & Brauner, 2008; Chromek et al., 2006; Nielsen et al., 2014; Spencer et al., 2012; Spencer et al., 2013). Remarkably, increased cathelicidin expression and LL-37 secretion is triggered a few minutes after bacteria encounter uroepithelial cells. This suggested role for AMPs in UTI immune defense is consistent with reports that UPEC strains are generally more resistant to AMPs than commensal *E. coli* strains that do not colonize the urogenital tract (Chromek et al., 2006).

Gram-negative bacteria use several mechanisms to resist killing by AMPs, including capsules, efflux pumps, LPS modifications, and proteases (Gruenheid & Le Moual, 2012). Omptin proteases are found in the Gram-negative outer bacterial membrane and have a conserved active site with features of both aspartate and serine proteases (Kramer et al., 2001; Vandeputte-Rutten et al., 2001). With their active sites facing the extracellular environment, omptins contribute to virulence by cleaving a variety of proteins and peptides (Haiko, Suomalainen, Ojala, Lahteenmaki, & Korhonen, 2009). Both substrate specificity and amino acid identity are used to classify omptins into Pla-like and OmpT-like subfamilies. Pla readily cleaves the proenzyme plasminogen into active plasmin to promote bacterial dissemination during both bubonic and pneumonic plague (Lathem, Price, Miller, & Goldman, 2007; Sodeinde et al., 1992; Zimbler, Schroeder, Eddy, & Lathem, 2015). OmpT rapidly cleaves and inactivates AMPs, including LL-37, protamine, and a synthetic peptide optimized to have maximum antibacterial activity called C18G (Brannon, Thomassin, Desloges, Gruenheid, & Le Moual, 2013; Stumpe, Schmid, Stephens, Georgiou, & Bakker, 1998; Thomassin, Brannon, Gibbs, Gruenheid, & Le Moual, 2012). OmpT-mediated AMP inactivation is thought to support host colonization by some pathogenic *E. coli* strains (Thomassin, Brannon, Gibbs, Gruenheid, & Le Moual, 2012). In addition to OmpT, two OmpT-like proteases have been described in *E. coli* strains (Kaufmann, Stierhof, & Henning, 1994; McPhee et al., 2014; Zhuge et al., 2018), these genes, called *ompP* and *arlC* (*ompTp*) encode proteins that have approximately 74% amino acid identity to OmpT. While the physiological substrates of OmpP and ArlC are unknown, OmpP has been shown to cleave the AMP protamine and ArlC is associated with AMP resistance (Hwang et al., 2007; McPhee et al., 2014).

The *ompT* gene is present in the genome of 85-97% of UPEC clinical isolates and is used in epidemiological studies to identify virulent UPEC strains, yet its function across clinical isolates remains unclear (Foxman, Zhang, Palin, Tallman, & Marrs, 1995). As OmpT and OmpT-like omptins play roles in resistance to host-produced AMPs, we hypothesized that high OmpT protease activity-levels contribute to UPEC colonization during symptomatic UTIs. To test this hypothesis, we detected *ompT* and measured OmpT activity in a collection of 58 UPEC clinical isolates from groups of patients with infections of differing clinical severity (fecal, ABU, UTI [cystitis and pyelonephritis] and sepsis). Heterogeneous OmpT activity was observed and in some isolates high protease activity was correlated with the presence of an additional ompT-like gene called *arlC* (*ompTp*). The presence of two OmpT-like proteases in some UPEC isolates led us to compare the substrate specificity of the three *E. coli* omptins (OmpT, OmpP and ArlC). We found that OmpT, OmpP and ArlC all cleave AMPs, although cleavage efficiency of different AMP-types varied. Our results suggest that the presence of multiple omptins allows UPEC to cleave at least two major subsets of AMPs encountered during infection.

## 2 Material and Methods

### 2.1 Bacterial Strains and Growth Conditions

58 ExPEC isolates originating from patients diagnosed with extraintestinal infections or from the urine or stool of healthy individuals were obtained from the Manges collection. Included isolates were randomly selected from the *E. coli* category to ensure they were representative. Isolates were divided into 4 groups based on disease type. Fecal isolates (*n* = 12) were recovered from the feces of healthy subjects in Québec Canada (2009-2010), ABU isolates (*n* = 10) were from patients with asymptomatic bacteriuria in California USA (2005-2006) (Manges, Johnson, & Riley, 2004), UTI isolates (*n* = 24) were recovered from patients with cystitis in Québec Canada (2005-2007) (Manges, Tabor, Tellis, Vincent, & Tellier, 2008) and cystitis or pyelonephritis in California USA (1999-2000) (Manges, Dietrich, & Riley, 2004), and sepsis isolates (*n* = 12) were from patients with sepsis in California USA (2001-2003) (Manges, Perdreau-Remington, Solberg, & Riley, 2006). Bacterial strains used in this study are listed in Table 1. Bacteria were routinely cultured in lysogeny broth (LB; 10% (w/v) tryptone, 5% (w/v) yeast extract, 10% (w/v) NaCl) or in N-minimal medium (50 mM Bis-Tris, 5 mM KCl, 7.5 mM (NH_4_)_2_SO_4_ 0.5 mM K_2_SO_4_, 0.5 mM KH_2_PO_4_, 0.1% casamino acids) adjusted to pH 7.5, supplemented with 1.4% glucose and 1 mM MgCl_2_ (UPEC isolates) or with 0.5% glucose and 1 mM MgCl_2_ (all other strains). Bacteria were cultured at 37°C with aeration (220 rpm).

**Table 1.**
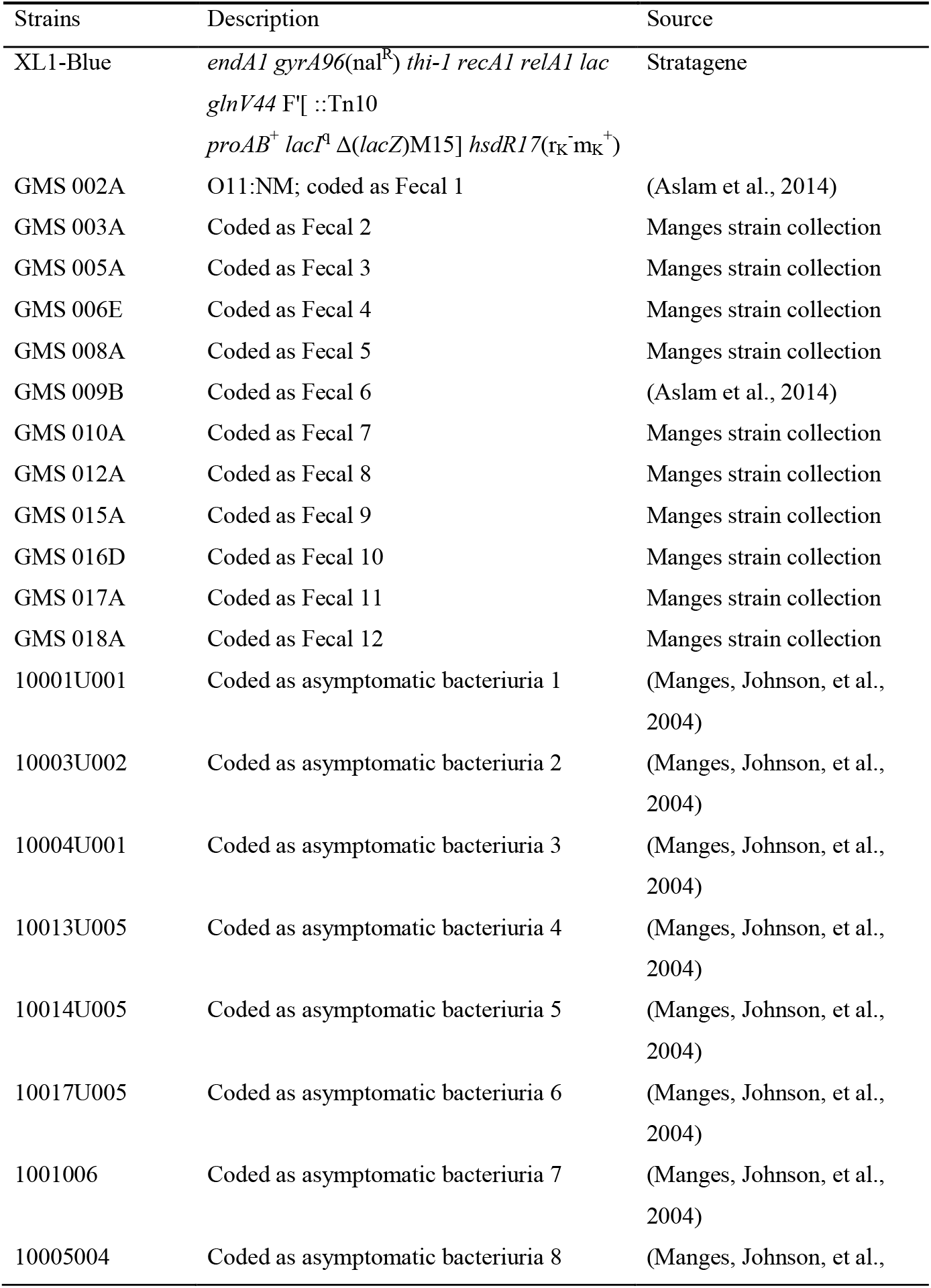
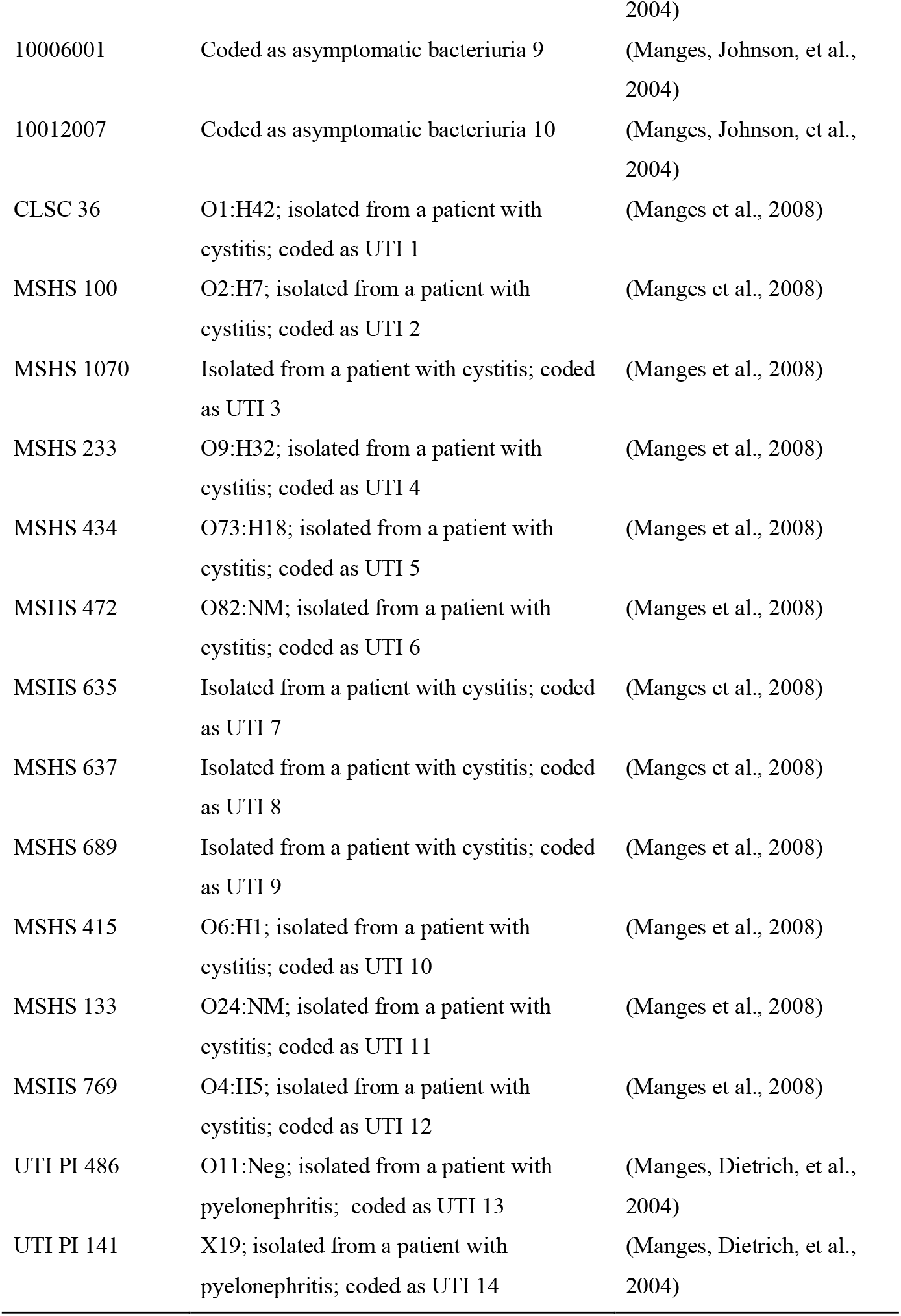
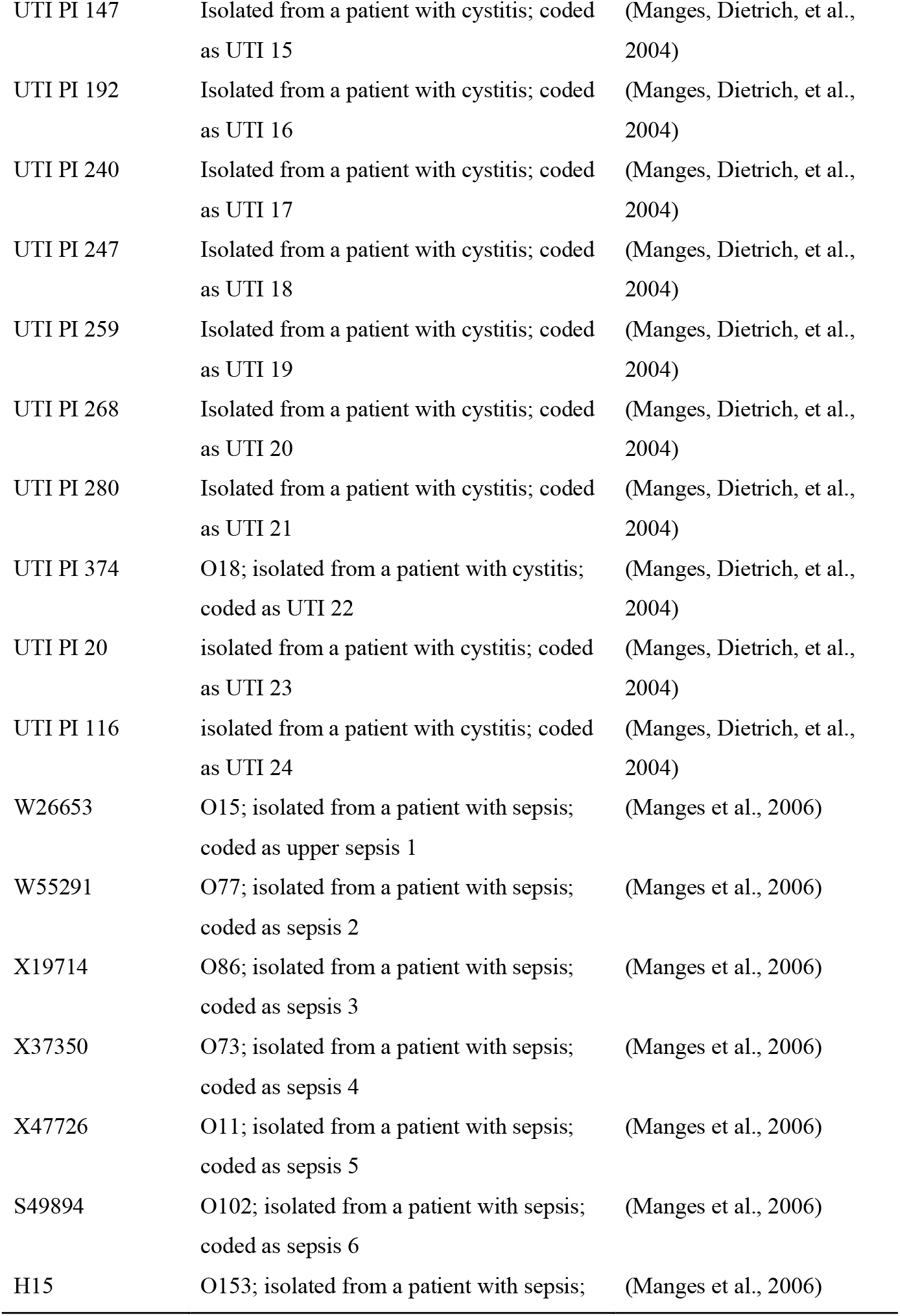
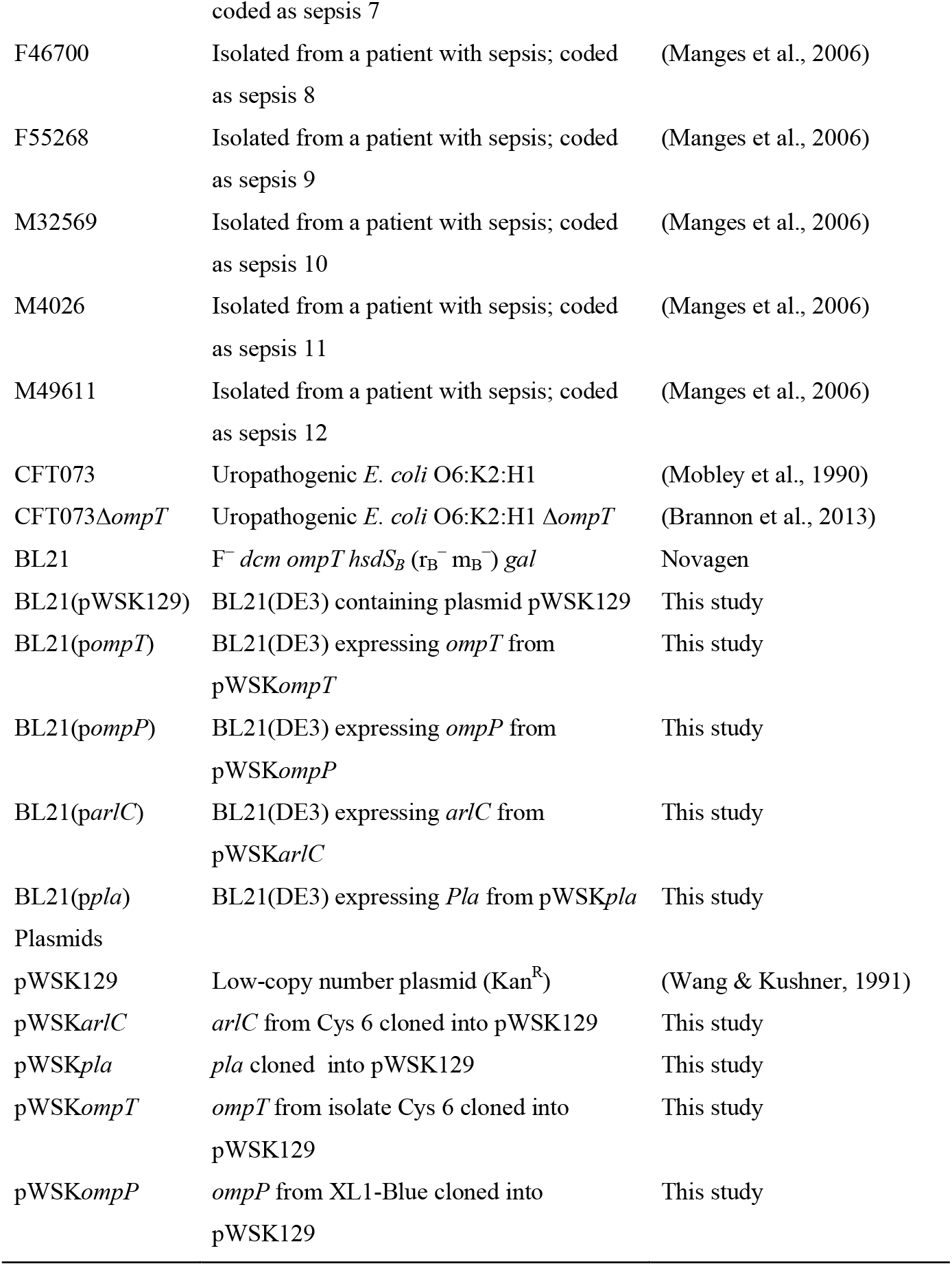
Strains and plasmids used in this study

### 2.2 Multiplex PCR of UPEC Virulence Genes

Total DNA (genomic and large-plasmid DNA) was isolated using the Puregene Yeast/Bact. kit (Qiagen). Phylogenetic groups were determined as described in (Clermont, Bonacorsi, & Bingen, 2000), using primer pairs listed in Table 2. To detect virulence genes present in the isolates, primer sequences were obtained from previous studies (Johnson & Stell, 2000) or designed *de novo* for this study (Table 2). Three multiplex PCR experiments were performed as follows: pool 1: *hylA* (1177 bp), *papAH* (720 bp), *fimH* (508 bp), *kspMTIII* (392 bp), and *papEF* (336 bp); pool 2; *papC* (200 bp), *sfaS* (240 bp), *cnf1* (498 bp), *fyuA* (880 bp), *iutA* (300 bp), *kpsMTII* (272 bp); pool 3: *arlC* (852 bp), *ompT* (670 bp) and *fimH* (508 bp); *ompP* (648 bp).

**Table 2.**
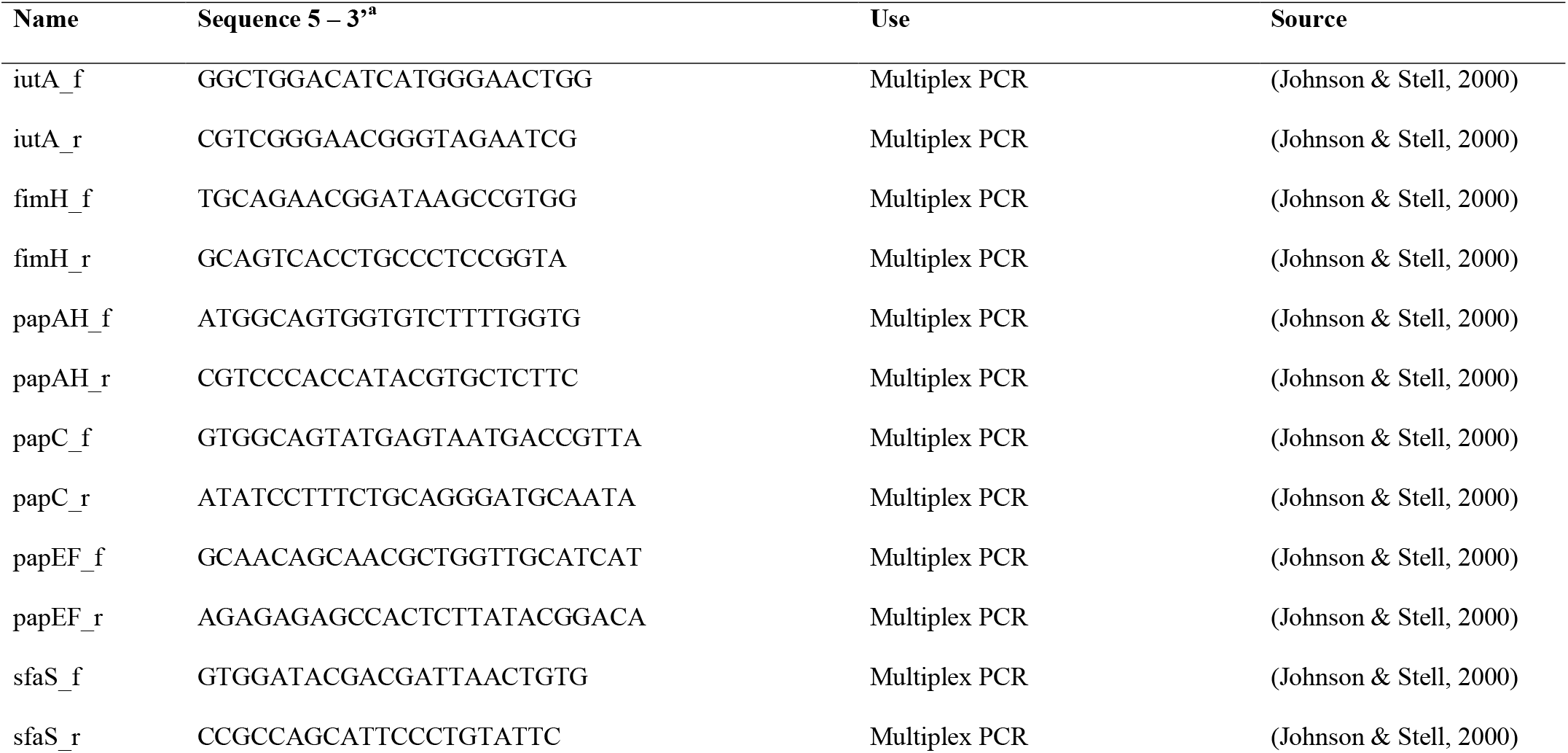
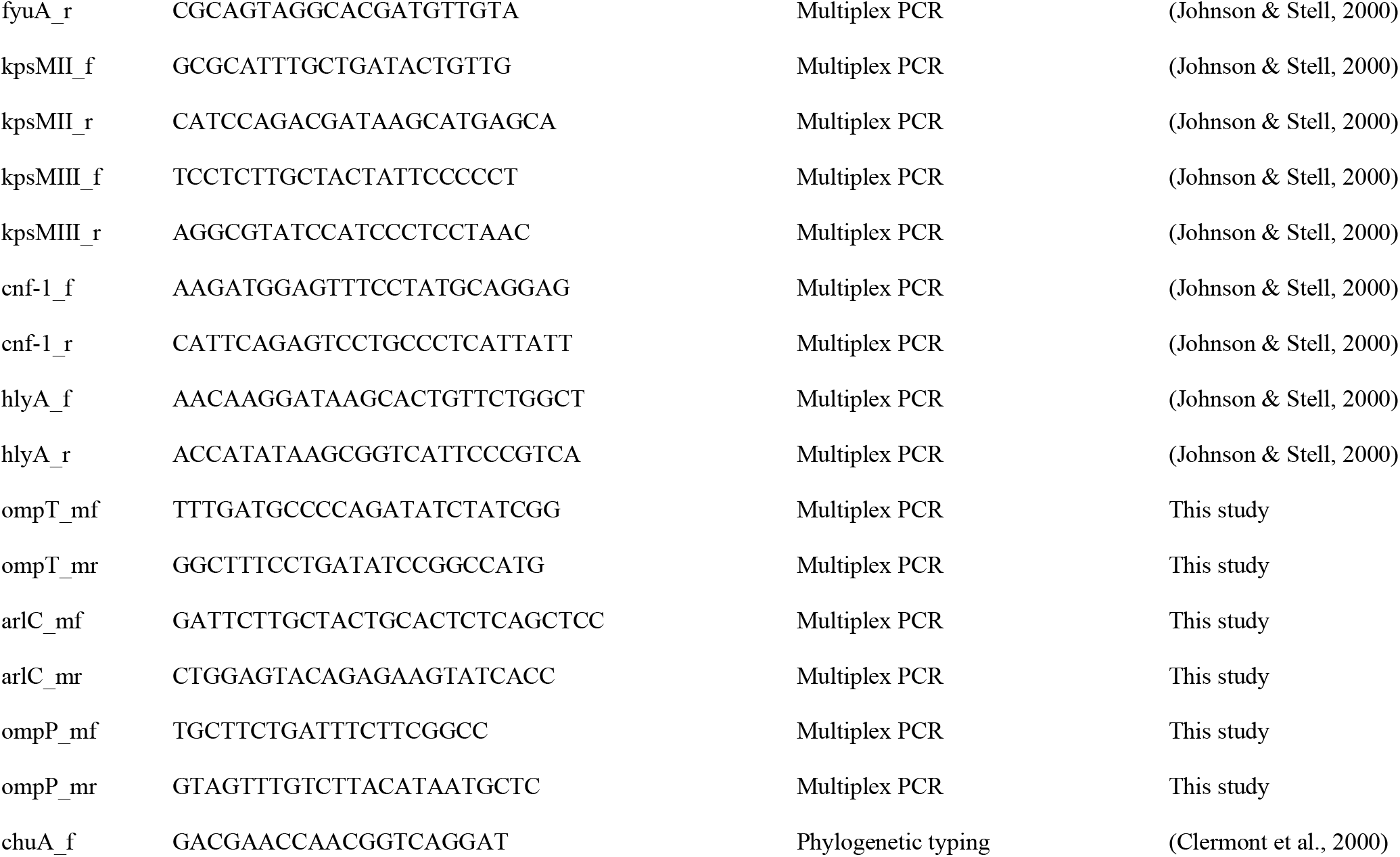
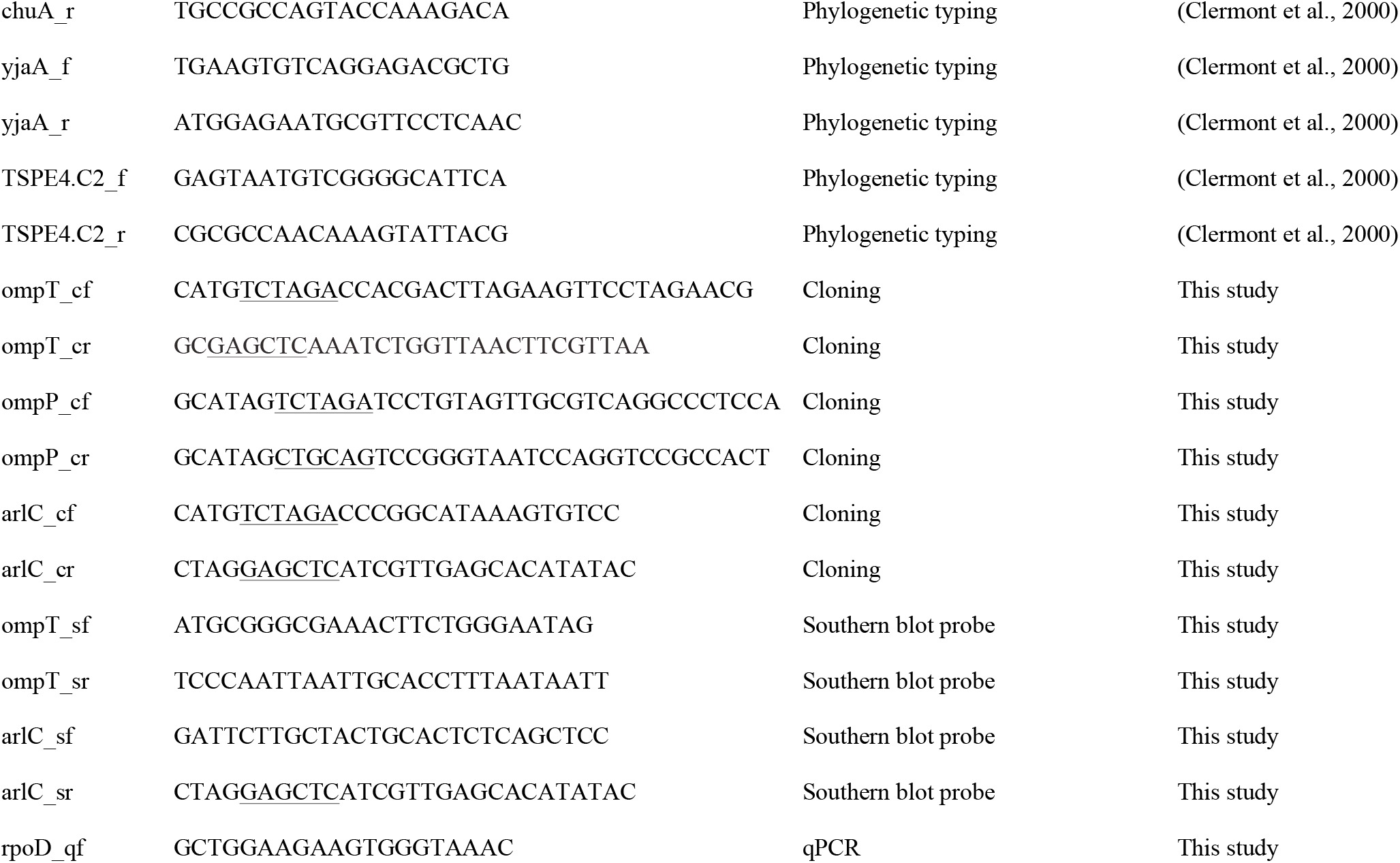
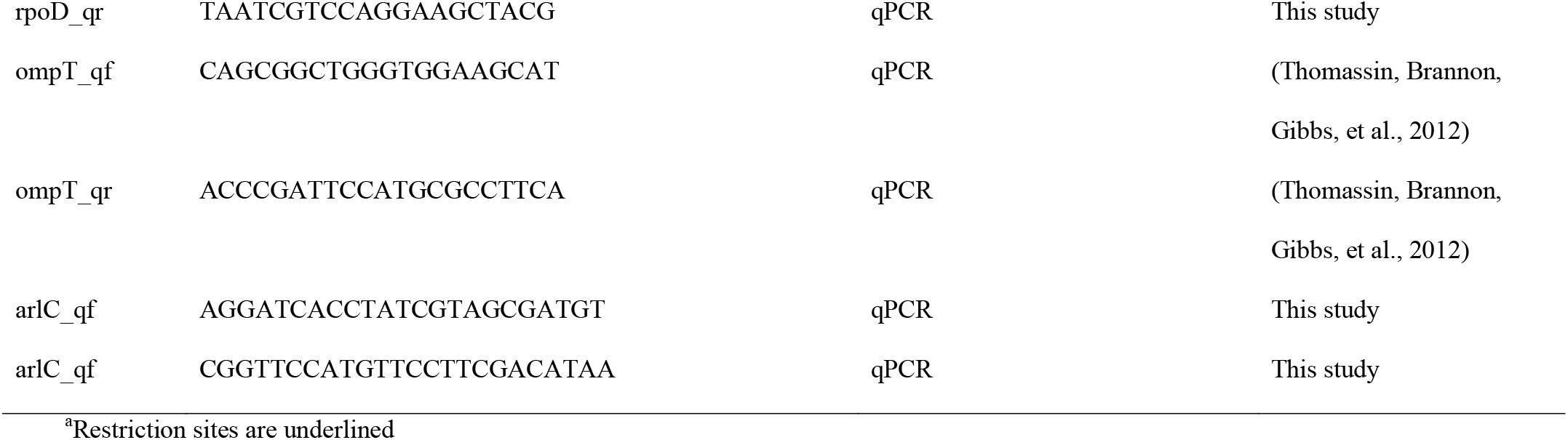
Primers used in this study

### 2.3 Fluorescence Resonance Energy Transfer (FRET) Activity Assay

The FRET substrate containing a dibasic motif (RK) in its center (2Abz-SLGRKIQI-K(Dnp)-NH2) was purchased from Anachem. Bacteria were grown in N-minimal medium to mid-exponential phase and normalized to an OD_595nm_ of 0.5. Bacterial cells were pelleted and resuspended in phosphate-buffered saline (PBS). Bacteria (~ 2.25 x 10^7^ CFU in 75 μL) were mixed in a 96-well plate with 75 μL of the FRET substrate (final concentration 3 μM). Fluorescence (λ Ex 325 nm, λ Em 430 nm) was monitored for 1 h at 25°C using a Biotek FLx 800 plate reader. Data were normalized by subtracting the background fluorescence of the FRET substrate in PBS.

### 2.4 Plasmid construction

The *ompT* and *arlC* genes were PCR-amplified from DNA isolated from the UPEC UTI clinical isolate 6, also called cystitis 6, using their respective primer pairs ompT_cf/ompT_cr and arlC_cf/arlC_cr (Table 2). PCR fragments were treated with XbaI and SacI and ligated into plasmid pWSK129 treated with the same enzymes, generating plasmids pWSK*ompT* and pWSK*arlC* (Table 1). The *ompP* gene was PCR-amplified from XL1-Blue DNA using primer pair ompP_cf/ompP_cr. PCR-products were treated with XbaI and PstI and ligated into pWSK129 treated with the same enzymes to generate plasmid pWSK*ompP*. The *pla* gene under control of the *croP* promoter was subcloned from pY*Cpla* (Brannon, Burk, et al., 2015) using XbaI and SacI and ligated into pWSK129 previously treated with the same enzymes, generating pWSK*pla*.

### 2.5 Southern Blotting

Total DNA was isolated and treated with EcoRV. Southern blotting and hybridization were performed as previously described (Taylor, Ouimet, Wargachuk, & Marczynski, 2011) using Hybond-XL membranes. Probes for *ompT* and *arlC* were PCR-generated using primer pairs ompT_sf/ompT_sr and arlC_sf/arlC_sr, respectively (Table 2). Probes were radiolabelled with dATP [α-32P] using the RadPrime kit (Invitrogen). The pWSK*arlC* plasmid was used as the positive control for the *arlC* probe.

### 2.6 Quantitative RT-PCR

Quantitative RT-PCR (qPCR) was performed as previously described (Thomassin, Brannon, Gibbs, et al., 2012). Briefly, bacterial strains were grown to an OD_595nm_ of 0.5 in N-minimal medium. Total RNA was isolated using TRIzol reagents (Invitrogen) and treated with TURBO DNase I (Ambion) to remove residual DNA. The absence of DNA was confirmed by qPCR using the primer pair rpoD_qf/rpoD_qr. RNA (100 ng) was reverse transcribed using Superscript II (Invitrogen) with 0.5 μg of random hexamer primers. A reaction mixture without Superscript II was also included and was used as the negative control. qPCR reactions were performed in a Rotor-Gene 3000 thermal cycler (Corbett Research) using the Maxima SYBR Green qPCR kit (Thermo Scientific), according to the manufacturer’s instructions. Primers used are listed in Table 2. The relative expression levels were calculated by normalizing the threshold cycle (*C*_T_) of *ompT* and *arlC* transcripts to the *C*_T_ of *rpoD* using the 2^−Δ*C*T^ method (Livak & Schmittgen, 2001).

### 2.7 Whole genome sequencing

Sequencing was performed on a PacBio platform (Pacific Biosciences). Genomic DNA samples were purified using the Gentra® Puregene® kit (Qiagen) and sheared to 20 kb using g-tubes (Covaris). Libraries were prepared using the template preparation kit from Pacific Biosciences. A single SMRT cell was sequenced to generate data sets including unique subreads with a minimum length of 3 kb. Genome assemblies of sequence reads were generated using a combination of HGAP/Celera/Quiver following Pacific Biosciences recommendations. The complete chromosome and plasmid sequences were submitted to GenBank.

### 2.8 Preparation of whole-cell lysates and outer-membrane fractions

Bacteria were grown in N-minimal medium until mid-exponential phase and normalized to an OD_595nm_ of 0.5. For whole-cell lysate samples, bacterial cells were pelleted and resuspended in 1/10 volume of 2X ESB (Thomas et al., 2005). Outer-membrane fractions were isolated as follows: bacterial cultures were centrifuged at 2500 rpm for 10 min and pellets were resuspended in 1.5 mL low-salt buffer (100 mM NaPi buffer [pH 7], 5 mM EDTA and 10% glycerol). Samples were supplemented with 10 uL PMSF and sonicated. Samples were then centrifuged at 5500 rpm for 10 min. Supernatants were collected and centrifuged at 65000 rpm for 30 min at 4°C. Pellets were resuspended in 2 mL sarcosyl buffer (10 mM Tris [pH 7.5], 5 mM MgCl_2_ and 2% sarcosyl) and incubated for 30 minutes at 10°C. Samples were then centrifuged for 60 min at 45000 rpm and the pellet containing outer membranes was resuspended in buffer (20 mM Tris-HCl pH 7.5 and 10% glycerol). Outer membrane samples were combined 1:1 with 2X ESB and boiled for 10 minutes prior to loading samples on an SDS-PAGE gel.

### 2.9 Western blotting

Whole-cell lysate and outer-membrane fractions were resolved on a 10% SDS-PAGE gel and transferred to a polyvinylidene fluoride membrane. Membranes were blocked for 1 h in Tris-buffered saline (TBS) supplemented with 5% skim milk and incubated overnight with the polyclonal anti-CroP antibody (Thomassin, Brannon, Gibbs, et al., 2012). Membranes were washed extensively with TBS and incubated for 1 h with a goat anti-rabbit secondary antibody conjugated with HRP. Membranes were washed and developed using chemiluminescent HRP substrate.

### 2.10 Plasminogen activation assay

Bacteria were grown in N-minimal medium to mid-exponential phase and normalized to an OD_595nm_ of 0.5. Bacterial cells were pelleted and resuspended in ½ volume of phosphate-buffered saline (PBS; final 6 x10^8^ CFU/mL). In a 96 well plate, 178 μL of bacteria and 20 μL of 45 mM VLKpNA (Sigma Aldrich) were combined. Baseline assays were performed at OD_405nm_. After 5 min, 4 μg of plasminogen substrate was added and absorbance (405nm) was measured every 10 min for 400 minutes at 37 °C with agitation before every reading.

### 2.11 Proteolytic cleavage of AMPs

Bacteria were grown in N-minimal medium to mid-exponential phase, washed and normalized to an OD_595nm_ of 0.5 in PBS. Aliquots of bacteria (10^7^ CFU) were combined 1:4 (v/v) with 2 μg/μL LL-37, mCRAMP, C18G or Magainin II (BioChemia) or 1 μg/μL RNase 7 and incubated at room temperature for various time points. Bacteria were separated from peptide cleavage products by centrifugation and supernatants were combined 1:1 with 2X ESB then boiled and frozen at −20°C. Peptide cleavage products were resolved on 10-20% Tris-Tricine gels (BioRad) and RNase 7 samples were resolved on 20% SDS-PAGE gels.

Peptides were fixed in the gel by incubation in 20% (v/v) glutaraldehyde for 30 min; gels were rinsed with water and peptides stained for 1h with Coomassie blue G-250 stain. Gels were destained in 20% (v/v) acetic acid.

### 2.12 Circular dichroism spectroscopy

Experiments were performed on a Jasco J-810 spectropolarimeter (Easton, MD). AMPs (200 μg/ml in PBS) were placed in a quartz cuvette with a path length of 0.1 cm and spectra were recorded from 260 to 195 nm. Samples were scanned three times at 20°C using a bandwidth of 1 nm, a time response of 2 sec and a scan rate of 100 nm/min. Spectra were corrected by subtracting the background spectrum of PBS and values were converted from ellipticity to mean residue ellipticity (MRE; degree × cm^2^ × dmol^-1^).

### 2.13 Statistical Analyses

Data were analyzed using GraphPad Prism software. Normality was verified using D’Agostino-Pearson normality test. Fisher’s Exact test was performed to compare incidence of virulence genes within severity groups of UPEC clinical isolates. FRET activity was assessed using a two way ANOVA with Tukey’s post test. *P* value ≤ .05 being significantly different.

## 3 RESULTS

### 3.1 Phylogenetic and virulence profile of UPEC isolates

UPEC isolates from patients with different disease severities were obtained from the Manges collection (Manges, Dietrich, et al., 2004; Manges et al., 2001; Manges, Johnson, et al., 2004; Manges et al., 2006; Manges et al., 2008). Although UPEC strains are heterogeneous, clinical isolates from UTIs predominantly belong to *E. coli* phylogenetic groups B2 and D (Johnson, Delavari, Kuskowski, & Stell, 2001). To confirm that our isolates are generally representative of UPEC clinical strains we determined the phylogenetic grouping of our 58 clinical isolates categorized into the fecal (n=12), ABU (n=10), UTI (cystitis and pyelonephritis; n=24), and sepsis (n=12) groups. Most isolates from the ABU and UTI groups associated with UTIs belong to the phylogenetic group B2 and, to a lesser extent, D (Table 3). In contrast, isolates from the sepsis group were predominantly from group D (Table 3). Finally, isolates from the fecal group had the most variable phylogenetic grouping with 5/12 isolates belonging to phyogenetic groups A and B1 (Table 3). Overall, this distribution is in agreement with previous reports showing that UPEC strains mainly belong to *E. coli* phylogenetic groups B2 and D (Johnson et al., 2001).

**Table 3.**
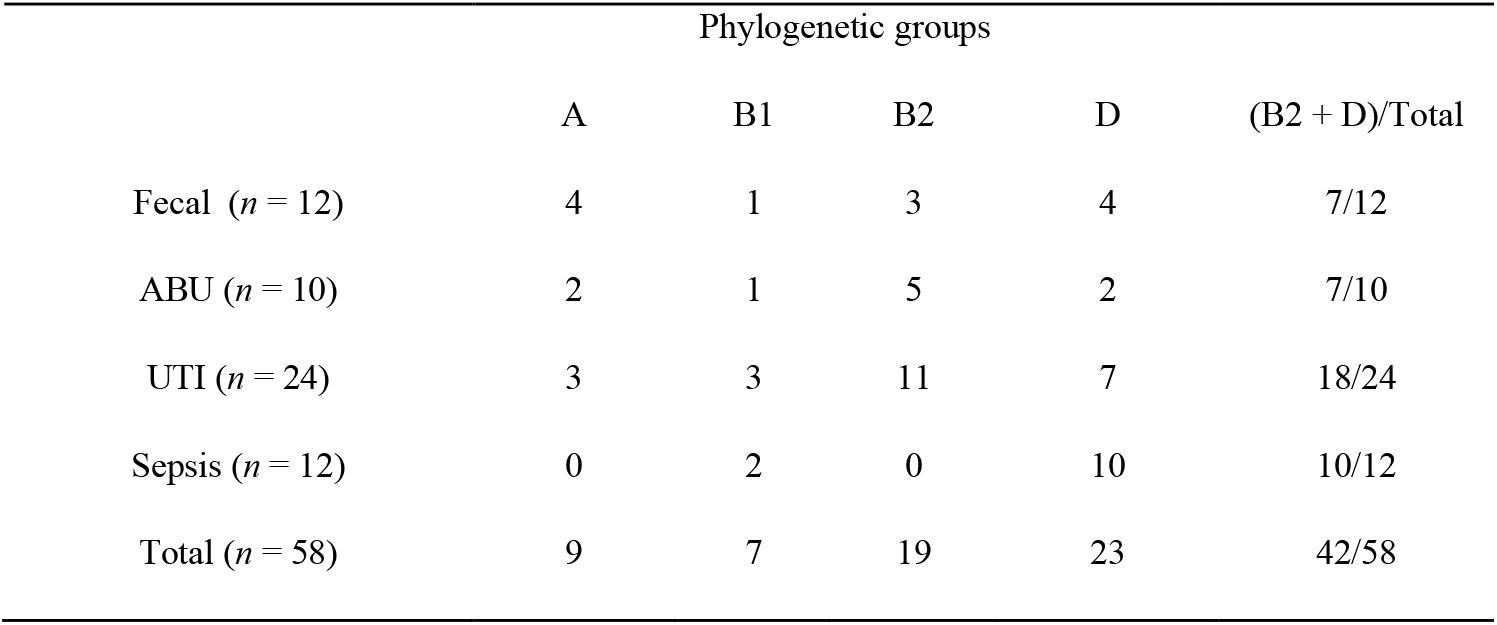
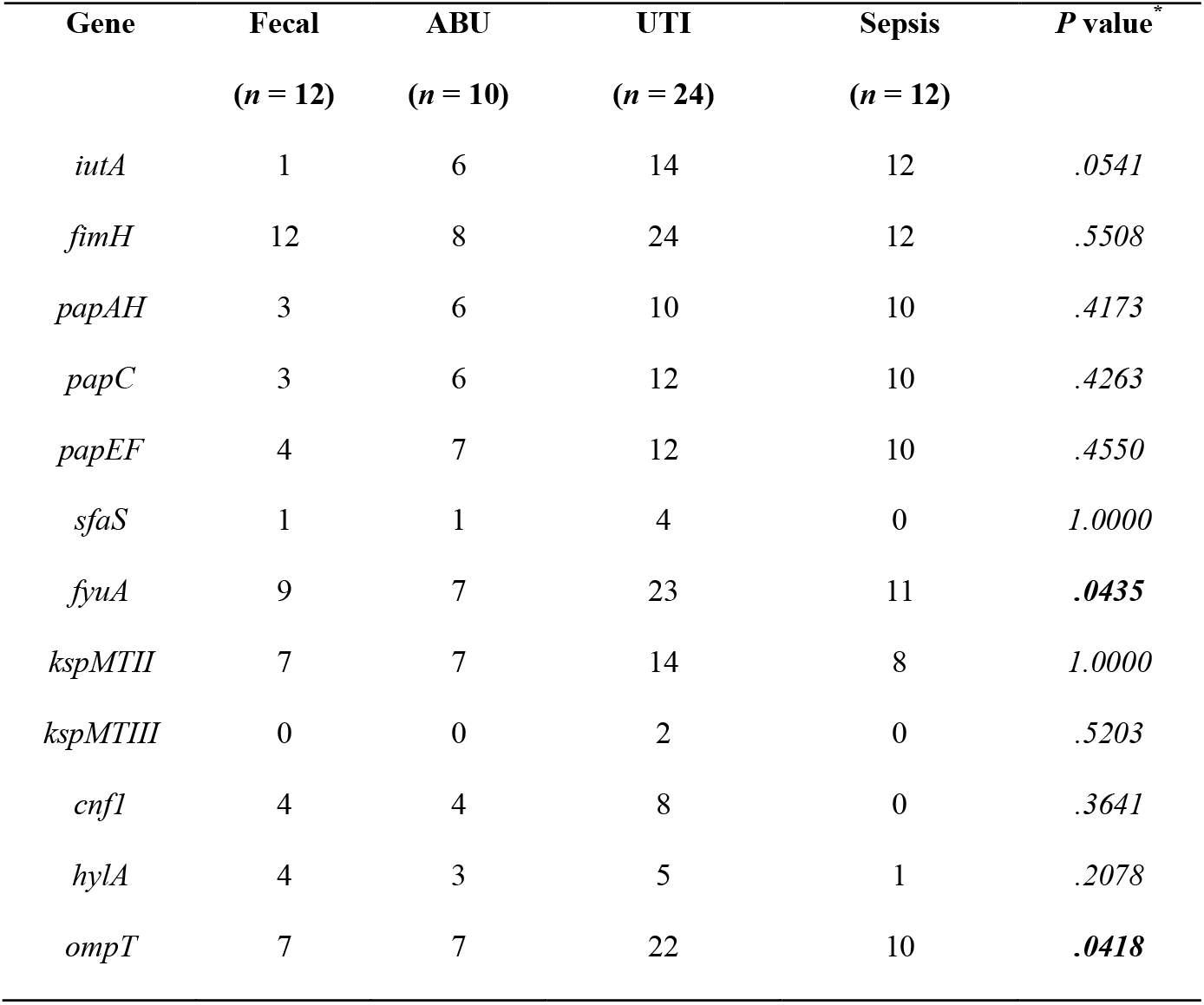
Phylogenetic distribution of UPEC clinical isolates.

The 58 isolates were further characterized using multiplex PCR to detect 12 recognized UPEC virulence genes (Table. 4). Our data showed variations consistent with previous studies reporting that UPEC is a heterogeneous pathotype (Marschall et al., 2012; Maynard et al., 2004; Norinder et al., 2012; Poey, Albini, Saona, & Lavina, 2012). The *fimH* gene, involved in UPEC adherence, was present in all but 2 ABU isolates (Table 4). There was a difference in the distribution of virulence genes *fyuA* and *ompT* for which the incidence was significantly higher in symptomatic (i.e UTI and sepsis) groups than asymptomatic (i.e. fecal and ABU) groups (Table 4). No other genes showed a significant difference in incidence between asymptomatic and symptomatic groups. In agreement with previous studies, we found that *ompT* is present in 89% of the UPEC isolates associated with symptomatic infections (Table 4).

### 3.2 Variability of omptin proteolytic activities among UPEC isolates

OmpT preferentially cleaves substrates between two consecutive basic residues (Dekker, Cox, Kramer, & Egmond, 2001; McCarter et al., 2004). Therefore, to assess OmpT proteolytic activity we measured cleavage of a FRET substrate (2Abz-SLGRKIQI-K(Dnp)-NH2) that contains a dibasic motif in its center (Brannon, Burk, et al., 2015; Brannon et al., 2013; McPhee et al., 2014; Thomassin, Brannon, Gibbs, et al., 2012). Cleavage of the substrate by the 58 UPEC isolates was monitored by measuring fluorescence emission over time and compared with substrate cleavage by the previously characterized reference UPEC strain CFT073 (Brannon et al., 2013). As shown in Fig. 1A, omptin activity of the isolates was heterogeneous between groups. Isolates for which the *ompT* gene was not detected by PCR showed basal activity levels (red triangles in Fig. 1A), whereas isolates harbouring the *ompT* gene showed a wide range of omptin activity. The omptin activity of the isolates of the fecal group was significantly lower than that of the 2 symptomatic groups (UTI and sepsis) (Fig. 1A). The mean activity of the isolates from the fecal group (0.75 ± 0.5) was lower than that of strain CFT073. In contrast, the activity means of the symptomatic groups (1.54 ± 0.66 and 1.71 ± 0.66) were higher than that of CFT073. Extensive variability in omptin activity was also observed within groups (Fig. 1A). The UTI group exibited the most heterogeneous omptin activity and some isolates from the UTI group had 3-fold higher omptin activity than CFT073. Together, these results indicate that omptin activity is variable among fecal and UPEC clinical isolates.

**Figure 1.**
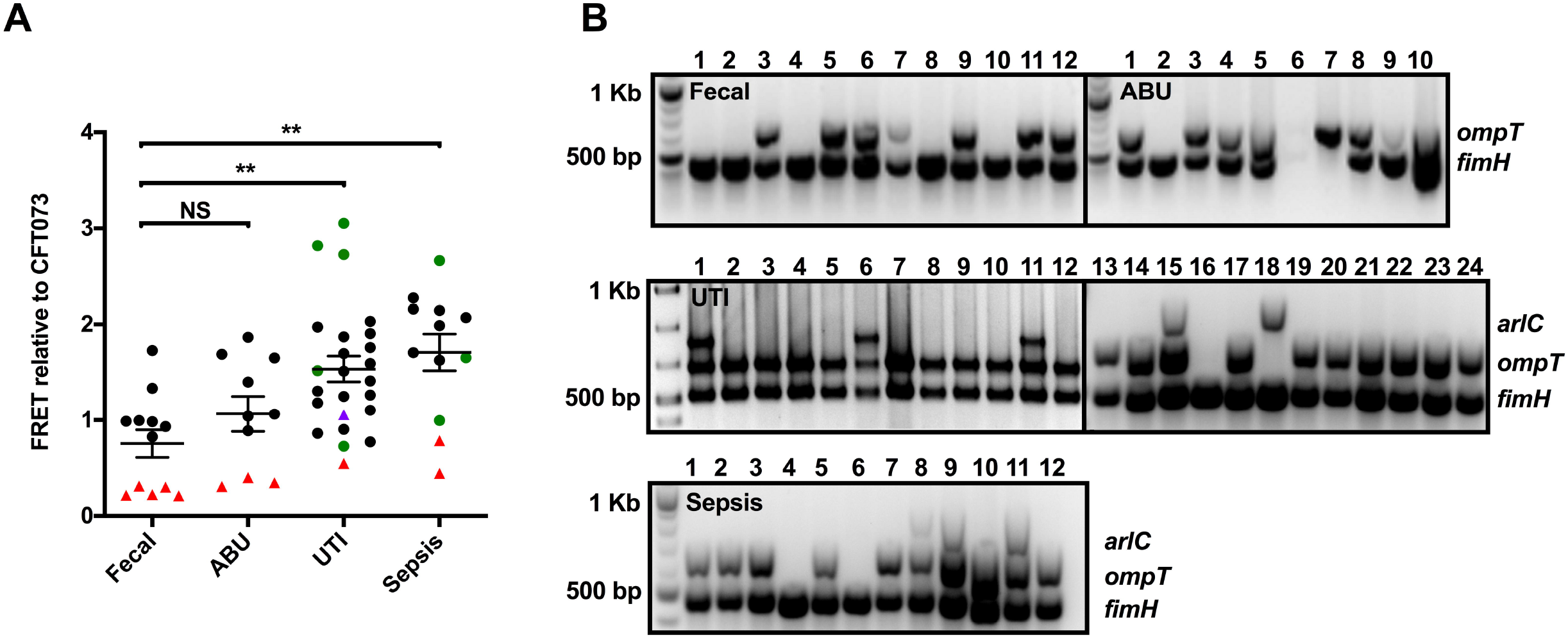
Omptin protease activity and distribution in clinical isolates. (A) Omptin activity was determined by monitoring fluorescence, indicative of FRET substrate cleavage, for 60 minutes. Data points indicate mean fold change in fluorescence of each isolate over the mean fold change in fluorescence of reference UPEC strain 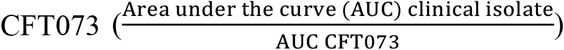 from triplicate samples. Bars represent mean ± SD fold change in fluorescence for each group. Bacteria that contain the *ompT* gene are indicated by circles and those that do not contain *ompT* are indicated by triangles. Indicated in green or purple are isolates that contain *arlC*. Statistical analysis was performed by one-way ANOVA followed by Tukey’s *post hoc* test using GraphPad Prism software (NS, not significant; *, *P* ≤ 0.05; **, *P* ≤ 0.01). (B) Multiplex PCR of *arlC* (852 bp), *ompT* (670 bp) and *fimH* (508 bp) from each of the clinical isolates. Amplification of *fimH* was used as a positive control. Numbers indicate isolate number for each group. Data are representative of at least three independent experiments.

### 3.3 OmpT-like proteases in UPEC

In addition to the chromosomally-encoded *ompT* gene, plasmid-borne *ompT*-like genes *ompP* and *arlC* are present in several *E. coli* strains (Kaufmann et al., 1994; McPhee et al., 2014; Zhuge et al., 2018). These OmpT-like proteins are approximately 74% identical to OmpT. To determine whether the presence of ompT-like genes in some isolates may account for the heterogeneity of OmpT activity observed in Fig. 1A, multiplex-PCR screens were performed to detect *ompT, ompP* and *arlC*. The *ompP* gene was not detected in any of the isolates (data not shown). In contrast, the *arlC* gene was present in 8 of the 58 isolates (Fig. 1B). Strikingly, *arlC* was only present in symptomatic isolates, which was statistcially significant according to a Fisher’s exact test (*P* = .0445). Most isolates harbouring the *arlC* gene also contained *ompT* and generally had higher proteolytic activity (green circles, Fig. 1A) than CFT073. This is consistent with the report that ArlC cleaves the FRET substrate (McPhee et al., 2014). Isolate 18 from the UTI group did not have *ompT* but harboured *arlC* (Fig. 1B); this isolate exibited moderate proteolytic activity (purple triangle in Fig. 1A). Together these data show that among commensal and clinical isolates, higher omptin activity is associated with symptomatic disease and isolates with the greatest omptin activity harbour both the *ompT* and *arlC* genes.

### 3.4 Variability of *ompT* and *arlC* expression among select UPEC cystitis isolates

To further understand omptin activity among UPEC isolates, we selected 12 isolates from the UTI group (Table 1) for further analysis because they have the most heterogeneous omptin activity. The presence of *ompT* genes in these isolates was confirmed by Southern blot analysis (Fig. 2A). This analysis also indicated that two *ompT* genes may be present in isolates 7, 8 and 11. Consistent with the multiplex PCR results, *arlC* was detected in UTI isolates 1, 6 and 11 (Fig. 2A). Next, qPCR was used to measure the expression levels of *ompT* and *arlC*. In agreement with our activity assay, *ompT*-levels were heterogeneous among these UTI isolates (Fig. 2B and 2C). Only three isolates (2, 10 and 11) had similar expression levels as the reference strain CFT073, whereas all other isolates had higher *ompT* expression levels than the reference strain. As expected from the multiplex-PCR screen and Southern blot, *arlC* expression was only detected in UTI 1, 6 and 11 isolates. UTI isolates 1 and 6, which showed the highest *ompT* and *arlC* expression levels also had the highest omptin activity-levels (Fig. 2C). Although both *ompT* and *arlC* are present in UTI isolate 11, they have low expression levels, which is consistent with the low omptin activity observed (Fig. 2C). These data indicate that heterogeneous omptin activity-levels are associated with both the presence and the different expression levels of the *ompT* and *arlC* genes.

**Figure 2.**
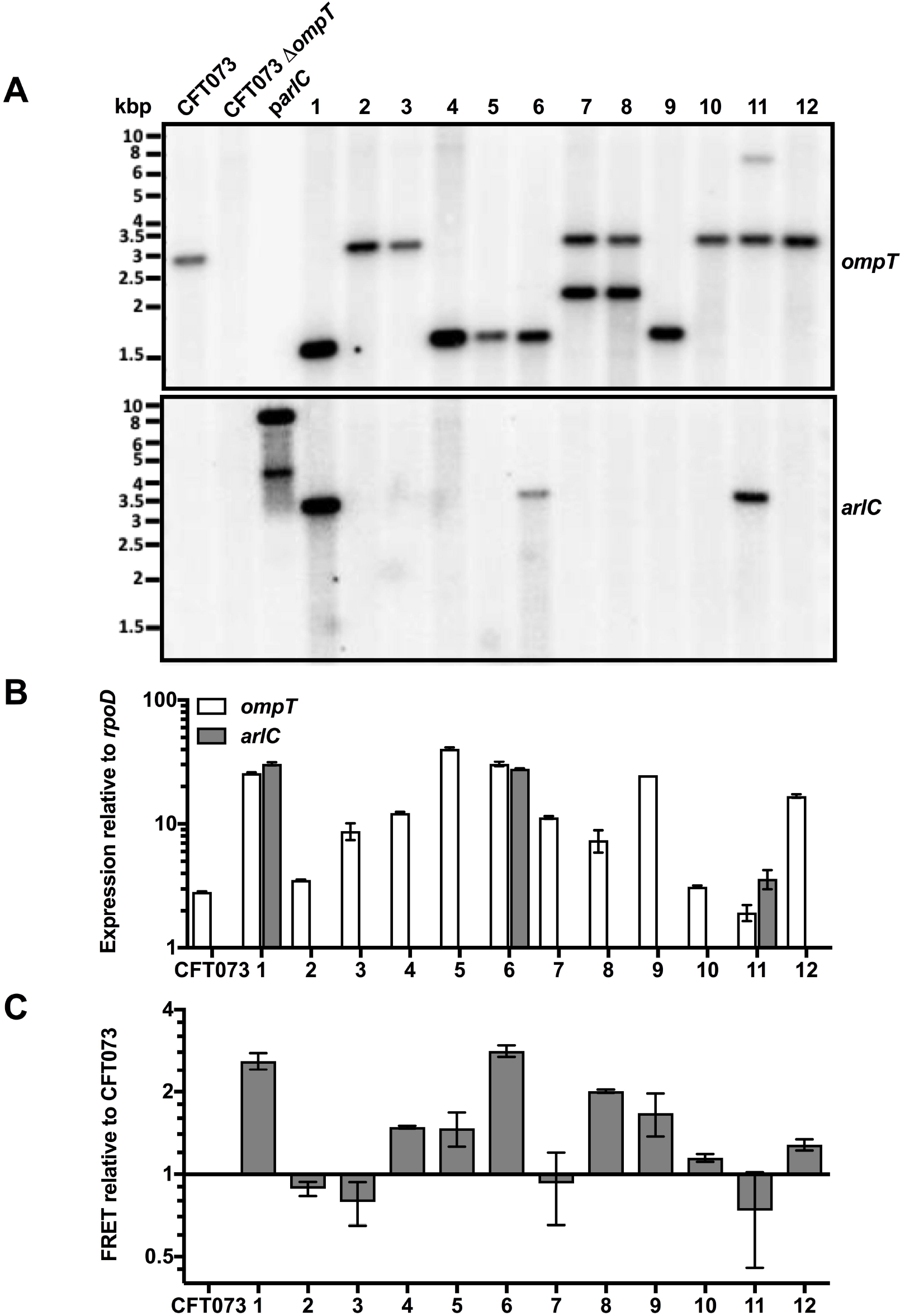
Presence and expression of *ompT* and *arlC* among select UTI isolates. (A) Southern blot of *ompT* and *arlC* from EcoRV-treated total DNA isolated from 12 cystitis-causing isolates, as well as control strains CFT073, CFT073Δ*ompT* and plasmid DNA from pWSK*arlC*. (B) Quantitative real time PCR (qRT-PCR) of *ompT* and *arlC* from 12 clinical isolates causing cystitis, as well as from reference strain CFT073. Shown is mean ± SD of *ompT* or *arlC* expression relative to *rpoD* calculated using the 2^−Δ*C*T^ method. Data are representative of 3 independent experiments. (C) Omptin activity of these cystitis clinical isolates was determined by monitoring cleavage of a synthetic FRET substrate for 60 minutes. Shown are mean ± SD change in fluorescence of each cystitis isolate over the change in fluorescence of reference stain 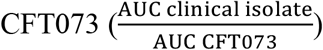. Data are representative of at least three independent experiments.

### 3.5 *arlC* is present on plasmids

To determine the genomic context of the *ompT* and *arlC* genes, isolates 1, 6 and 11 of the UTI group were sequenced on a PacBio platform. These isolates were then renamed cystitis 1, cystitis 6 and cystitis 11. Detailed descriptions of genomes and gene features are found in Appendix (Appendix Figs. 1AB, 2AB, 3). In all three isolates, *ompT* was located within the bacterial chromosome and *arlC* was part of large plasmids (79-200 kbp; Appendix Fig. 1A and 2A). In addition, the *ompT* gene was invariably located downstream of *nfrA* and *ybcH* (Fig 3A). Some differences were noted in the genomic context of *ompT* among the clinical isolates. In cystitis 1 and 6 the *envY* gene, encoding a transcriptional regulator of porin synthesis, is inserted between *ybcH* and *ompT* (182 bp downstream of *ybcH*, 512 bp upstream of *ompT*). The *appY* gene, encoding a transciptional activator, is located 249 bp downstream of the *ompT* gene in cystitis 1, whereas *ymcE*, encoding a putative cold shock gene, is located 186 bp downstream of *ompT* in cystitis 6. In cystitis 11 the *ompT* gene is located 657 bp downstream of *ybcH* and 272 bp upstream of *ybcY*; this is the same genomic context as that in UPEC strains CFT073, UTI89, 536, J96 ABU83972 and EPEC strain e2348/69, all of which were reported to have low omptin activity (Fig. 3A, (Brannon et al., 2013; Thomassin, Brannon, Gibbs, et al., 2012; Thomassin, Brannon, Kaiser, Gruenheid, & Le Moual, 2012)). For all isolates, the predicted amino acid sequence of ArlC is 100% identical to ArlC identified in adherent-invasive *E. coli* (AIEC) strain NRG857c (McPhee et al., 2014). Although the three plasmids harbouring *arlC* were different (Appendix Fig. 2A and B), *arlC* was present in all cases as part of pathogenicity island PI-6 previously reported to play a role in AMP resistance (Fig. 3B, McPhee et al., 2014).

**Figure 3.**
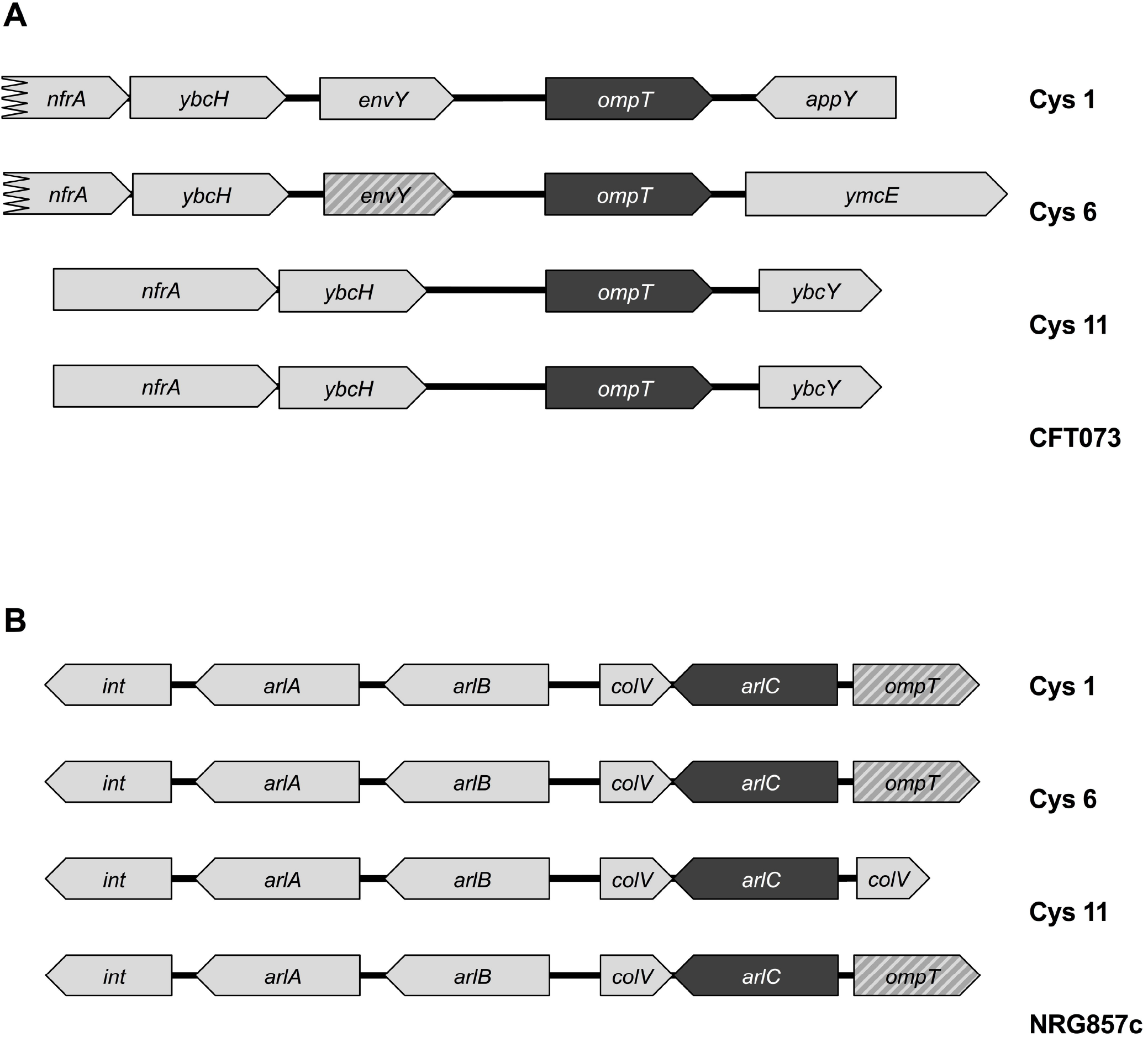
Genomic context of *arlC* and *ompT*. Schematic representation of the genomic contexts of the *ompT* (A) and *arlC* (B) genes in cystitis isolates 1, 6 and 11. Genomic contexts of *ompT* (A) and *arlC* (B) from respective reference strains CFT073 (A) and NRG857c (B) are included for comparison. Omptin genes are indicated in dark gray, light gray indicates genes located upstream and downstream of the omptin genes, stripes indicate pseudogenes and black lines indicate intergenic space.

### 3.6 Comparative analysis of OmpT, OmpP and ArlC

With the unexpected detection of *arlC* among the UPEC clinical isolates, we hypothesized that the presence of a second or even a third omptin protease within a single species may provide an advantage by expanding the potential range of substrates cleaved. Therefore, we sought to compare the substrate specificities of these proteases. As OmpT undergoes autocleavage during purification (Kramer, Zandwijken, Egmond, & Dekker, 2000; Vandeputte-Rutten et al., 2001) and mutagenesis of residues to stabilize the protein results in a significant decrease in FRET substrate cleavage ((Kramer et al., 2000); unpublished data Thomassin JL and Brannon JR) it was not possible to purify these proteases and directly compare their activities. Instead, we produced OmpT, OmpP and ArlC in *E. coli* BL21, a laboratory strain that lacks omptin proteases. To test their production and correct localization in BL21, omptin proteins were detected by western blot analysis from both whole cells and outer-membrane preparations (Fig. 4A). To determine if the proteases were active in BL21, FRET substrate cleavage was monitored over time. As expected, BL21 with empty vector did not cleave the FRET substrate, whereas the three omptins readily cleaved the FRET substrate (Fig. 4B). This demonstrates that when produced in BL21, ArlC, OmpP and OmpT are found in the outer membrane and are proteolytically active.

**Figure 4.**
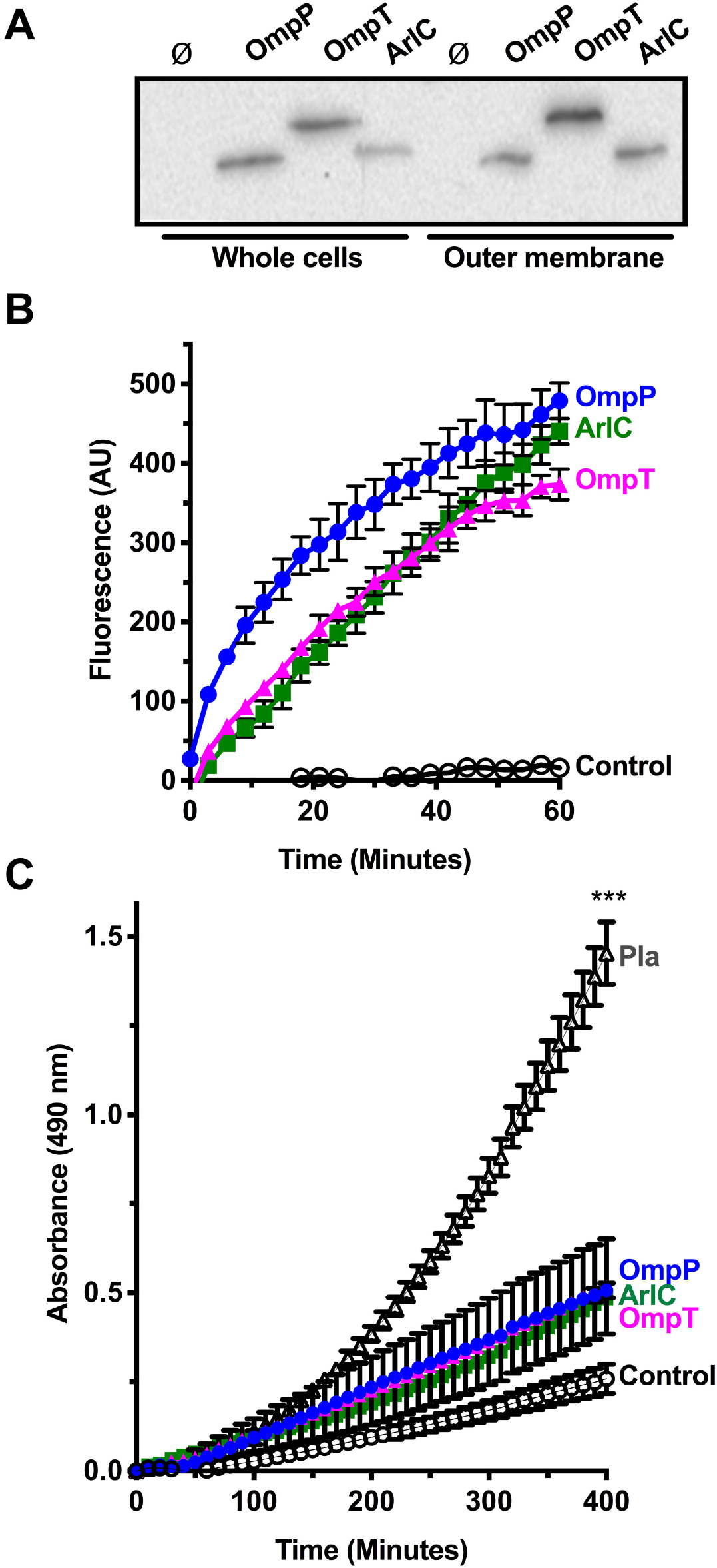
ArlC, OmpP and OmpT are functional in BL21. (A) BL21 containing empty vector (ø) or plasmids encoding *arlC, ompP* or *ompT* were grown until mid-log phase and normalized to OD_595_ 0.5. Proteins from whole cell preparations or isolated bacterial outer membranes were resolved by SDS-PAGE and transferred to a PVDF membrane. Omptins were detected by western blot using anti-CroP polyclonal antibodies. (B) A synthetic FRET peptide containing a dibasic motif (RK) was incubated with BL21 (open circles; control) or BL21 expressing *arlC* (filled squares; ArlC), *ompP* (filled circles; OmpP) or *ompT* (filled triangles; OmpT). Peptide cleavage, indicated by increased fluorescence, was monitored over time. Data show the mean ± SD from triplicate samples and are representative of at least three independent experiments. (C) Plasmin activation by ArlC, OmpP and OmpT. Glu-plasminogen and VLKpNA (plasmin substrate) were incubated with BL21 (open circles; control), BL21(p*pla*) (open triangles; Pla), BL21(p*arlC*) (filled squares; ArlC), BL21(p*ompP*) (filled circles; OmpP), or BL21(p*ompT*) (filled triangles; OmpT) strains. Absorbance at 405 nm was monitored over time. Data were normalized by subtracting initial absorbance from all values. Data represent mean ± SD and are representative of at least three independent experiments.

Omptin proteases are generally subdivided into OmpT-like or Pla-like subfamilies. These subfamilies differ in their ability to cleave plasminogen to activate it into active plasmin, with Pla-like omptins producing active plasmin more readily than OmpT-like omptins (Haiko et al., 2009; Kukkonen et al., 2001). To verify that the three omptin proteases belong in the OmpT-like subfamily, we tested their ability to cleave plasminogen into plasmin. Consistent with their presence in the outer membrane, all three omptins cleaved plasminogen to a greater extent than BL21 alone (Fig. 4C). There was no difference in their ability to activate plasminogen. Compared with the positive control, Pla produced in BL21, the *E. coli* omptins converted significantly less plasminogen into plasmin. These data are consistent with previous publications (Brannon, Burk, et al., 2015; Kukkonen et al., 2001; McPhee et al., 2014) and suggest that all three omptins found in *E. coli* belong to the OmpT-like subfamily of omptin proteases.

Omptin proteases belonging to the OmpT-like subfamily have been associated with AMP cleavage (Le Sage et al., 2009; Stumpe et al., 1998; Thomassin, Brannon, Gibbs, et al., 2012). Previous work has shown that OmpT from EPEC, EHEC, and UPEC cleave the human cathelicidin LL-37. Although ArlC was shown to play a role in AMP-resistance (McPhee et al., 2014), and OmpT and OmpP are reported to exhibit similar substrate specificities (Hwang et al., 2007; McCarter et al., 2004), their ability to cleave different AMPs has not been directly compared. Therefore, we investigated the ability of the *E. coli* omptins to cleave the synthetic cationic peptide C18G and various cathelicidins Magainin II (*Xenopus laevis*), mCRAMP (*Mus musculus*) and LL-37 (*Homo sapiens*). As expected, AMPs incubated with BL21 did not show any degradation or cleavage products, indicating that BL21 does not contain intrinsic proteases that cleave these AMPs (Fig 5A). OmpT cleaved all peptides by the first time point tested (2 min C18G; 15 min mCRAMP, Magainin II and LL-37; Fig. 5A). Similarly to OmpT, OmpP readily cleaved C18G and Magainin II within 2 and 30 min, respectively. In contrast, OmpP only cleaved small amounts of mCRAMP after 60 min and did not appear to cleave LL-37 (Fig. 5A). ArlC cleaved mCRAMP, C18G and Magainin II by the first time point tested (2 min C18G; 15 min mCRAMP and Magainin II), but only a small amount of LL-37 cleavage was observed after 60 minutes. Substrate properties, such as size and secondary structure are known to influence omptin activity (Brannon, Thomassin, Gruenheid, & Le Moual, 2015; Hritonenko & Stathopoulos, 2007). Given that all three proteases readily cleave the FRET substrate and C18G, the striking differences in ability to cleave Magainin II, mCRAMP and LL-37 are likely due to intrinsic differences between OmpT, OmpP and ArlC. Although all peptides tested contain sites with two consecutive basic residues, they range in size from 18-37 amino acids (Fig. 5B). OmpP cleaved smaller peptides such as C18G and Magainin II more readily than the larger mCRAMP and LL-37. As ArlC cleaved mCRAMP more rapidly than OmpT, peptide length might not be the limiting factor for this protease. Peptide secondary structure also influences omptin activity (Brannon, Thomassin, et al., 2015), therefore, we used circular dichroism spectroscopy to determine the secondary structure of these AMPs (Fig. 5C). Under our experimental conditions, only LL-37 is α-helical, while mCRAMP, C18G and Magainin II are unstructured (Fig. 5C). While peptide structure did not affect OmpT activity, ArlC did not appear to cleave the only α-helical AMP (Fig. 5BC). Together these findings suggest that OmpT, OmpP and ArlC have differences in substrate cleavage specificities.

**Figure 5.**
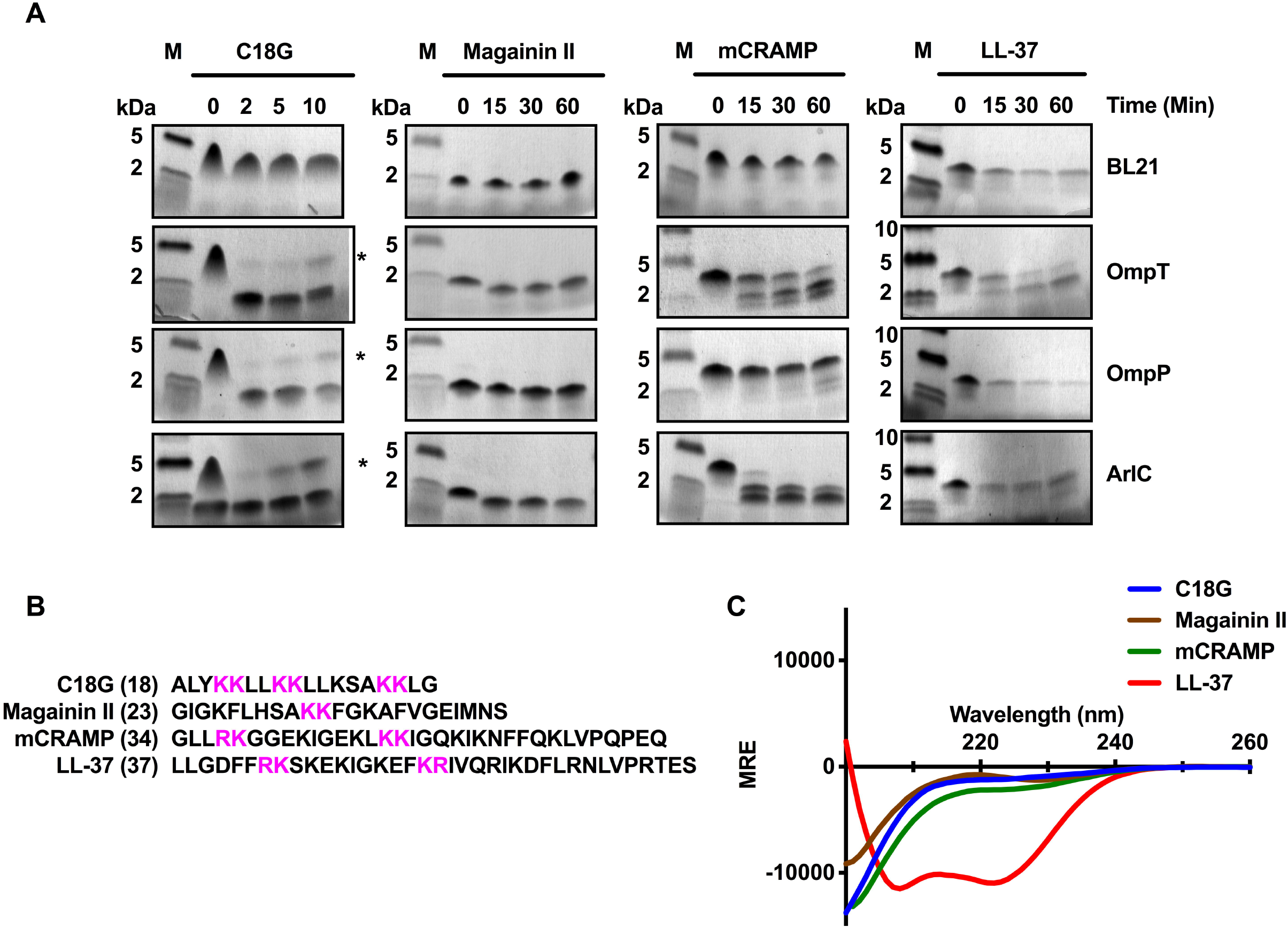
ArlC, OmpP and OmpT cleave cathelicidins. (A) AMP-cleavage assay. AMPs were incubated with BL21 alone or BL21 expressing *arlC, ompP* or *ompT* for the indicated times. Resulting AMP-cleavage products were separated by Tris-Tricine SDS-PAGE, fixed with glutaraldehyde and visualized by coomassie staining. M indicates molecular weight marker. Data are representative of three independent experiments. (B) Amino acid sequence of AMPs cleaved in (A) with dibasic motifs highlighted in magenta and sequence length indicated in parenthesis. (C) Far UV circular dichroism spectra (200-260 nm) of the indicated peptides measured in PBS. Data were normalized by subtracting spectra from PBS alone from the sample spectra. MRE indicates degree × cm^2^ × dmol^−1^.

We previously reported that disulfide bonds present in defensins render them resistant to OmpT-mediated proteolysis (Thomassin, Brannon, Kaiser, et al., 2012). Yet ArlC was shown to contribute to bacterial survival in the presence of human defensins (McPhee et al., 2014), suggesting that unlike OmpT, ArlC might cleave AMPs that are stabilized by disulfide bridges. RNase 7 contains four disulfide bridges, three dibasic sites (Fig. 6A) and is abundant in the urinary tract (Spencer et al., 2011; Spencer et al., 2013). The presence of dibasic sites suggests that RNase 7 might be an omptin substrate; therefore, we sought to investigate if there was a difference in omptin-mediated cleavage of this peptide. Under our experimental conditions, OmpT and OmpP did not cleave RNase 7 (Fig. 6B). After 60 min incubation with ArlC, an RNase 7 cleavage product appeared, with more cleavage product appearing after 90 min. While cleavage appears limited, ArlC was the only OmpT-like omptin able to cleave RNase 7. Taken together, these data indicate that ArlC, OmpP, and OmpT have different substrate specificities, suggesting that the presence of multiple omptin proteases in a single bacterial strain may enhance AMP-resistance by increasing the range of substrates cleaved.

**Figure 6.**
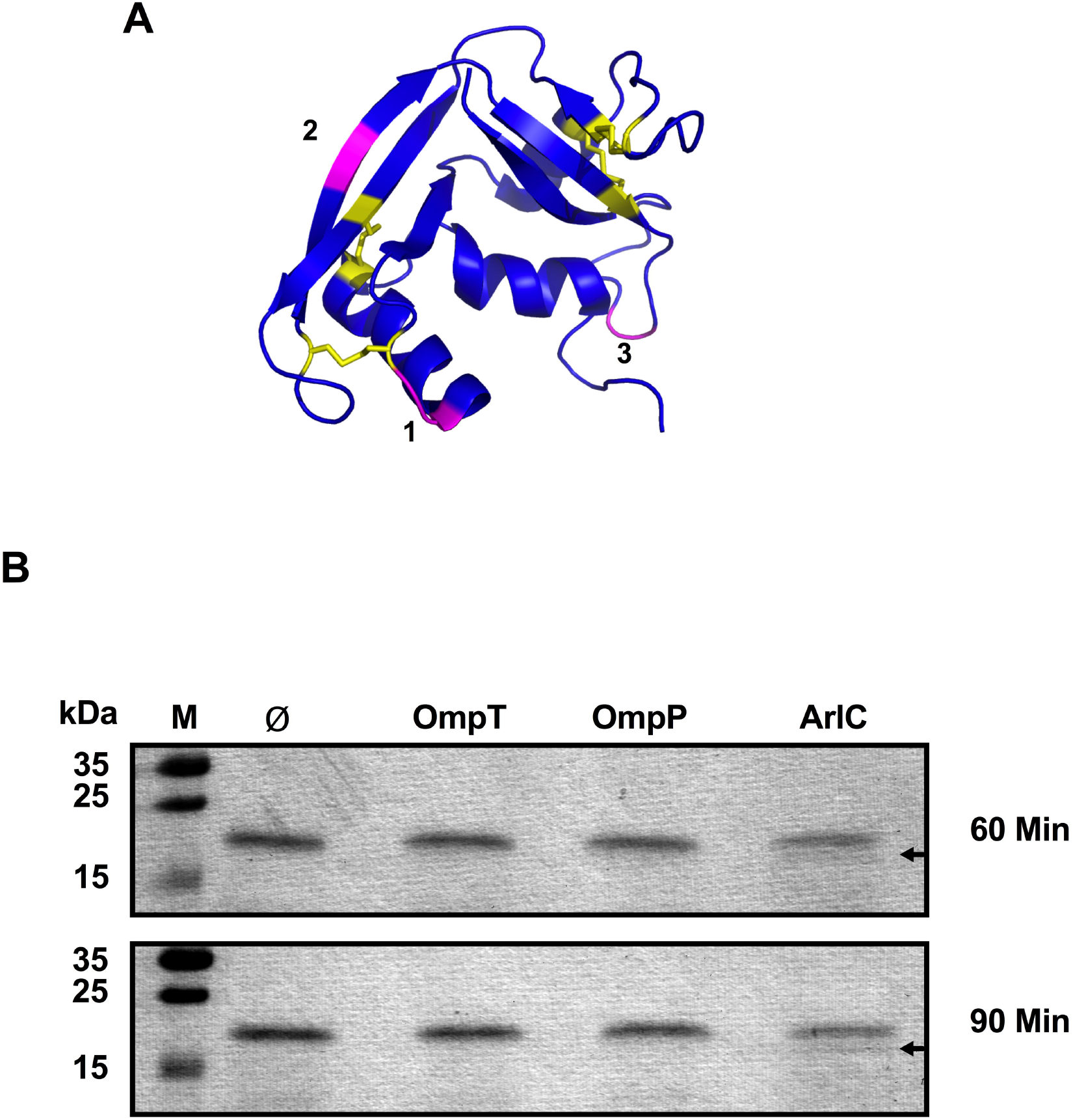
ArlC cleaves RNase 7. (A) Pymol generated image of RNase 7 (Refs; PDB 2hky), peptide backbone is shown in blue, cysteines and disulfide bridges are in yellow and dibasic sites are coloured magenta. Numbers correspond to the following the locations of the dibasic sites in the protein sequence: 1, residues 35 and 36; 2, residues 96 and 97; 3, residues 111 and 112. (B) Proteolytic cleavage of RNase 7. RNase 7 was incubated with BL21 containing empty vector (ø) or BL21 expressing *arlC, ompP* or *ompT* for 60 or 90 minutes. Cleavage products (arrows) were separated by SDS-PAGE and visualized by coomassie staining. M indicates molecular weight marker. Data are representative of three independent experiments.

## 4 DISCUSSION

Detection of specific genes, including *ompT*, is often used to characterize virulent clinical UPEC isolates (Johnson et al., 2001; Najafi et al., 2018). Previous studies have suggested that OmpT from the UPEC strain CFT073 is involved in adhesion, invasion and/or inactivation of AMPs (Brannon et al., 2013; He et al., 2015). While the presence of *ompT* is associated with virulent strains, its precise contribution remains unclear, as UPEC clinical isolates have highly variable genetic sequences (Schreiber et al., 2017). In addition, we previously observed large differences in OmpT protein activity due to differential *ompT* expression (Thomassin, Brannon, Gibbs, et al., 2012; Thomassin, Brannon, Kaiser, et al., 2012) suggesting that the presence of the *ompT* gene may not entirely correlate with its activity levels in different UPEC clinical isolates. In this study we hypothesized that OmpT activity correlates with increased disease severity among UPEC clinical isolates. To test this hypothesis, we systematically measured omptin activity in 58 *E. coli* isolates representing colonization and a range of clinical outcomes. Increased omptin activity was correlated with clinical UPEC strains isolated from patients with symptomatic UTIs (UTI and sepsis groups).

Omptin activity was heterogeneous among the clinical isolates, and could be related with differential *ompT* expression and the presence of a second OmpT-like protease, *arlC*. For example, a 20-fold difference in *ompT* expression was observed between isolates 5 and 11 of the UTI group (Fig. 2B). This finding is not unprecedented, since it was previously shown that *ompT* expression was 32-fold higher in EHEC than in EPEC (Thomassin, Brannon, Gibbs, et al., 2012). Differential *ompT* expression-levels in EHEC and EPEC were attributed to differences in distal promoter sequences found more than 150 bp upstream of the *ompT* start codon (Thomassin, Brannon, Gibbs, et al., 2012). An EPEC-like *ompT* distal promoter sequence and genomic context was also correlated with low OmpT activity in UPEC reference strains (Brannon et al., 2013). Therefore, it was not surprising that the EPEC-like promoter in cystitis (UTI) isolate 11 resulted in low *ompT* expression and OmpT activity. The insertion of *envY* in the intergenic space between *nfrA* and *ompT* correlated with the increased *ompT* expression and OmpT activity levels observed in cystitis (UTI) isolates 1 and 6 (Figs. 2BC and 3A). These data further suggest that variations in distal promoter sequences are responsible for differential *ompT* expression and, in turn, proteolytic activity observed. It is also possible that in addition to differences in the promoter regions, transcription factors or post-transcriptional factors regulating *ompT* expression are absent or differentially expressed in some isolates. Another explanation for heterogeneous omptin activity observed in this study can be attributed to the presence of a second plasmid-encoded omptin, *arlC*, in some isolates. The *arlC* gene was first identified as part of a large virulence plasmid of the AIEC strain NRG857c (McPhee et al., 2014). BLAST searches in the NCBI database revealed that *arlC* can also be found on plasmids harboured by various human ExPEC strains isolated from patients with meningitis and sepsis, as well as avian *E. coli* strains (Appendix Fig. 2B). While we did not detect *ompP* in our study, *ompP* is present in some UPEC strains that were collected and sequenced by the Broad Institute (“E.coli UTI Bacteremia initiative,” 2019). This opens the possibility that any combination of ompT-like omptin may be present in a given UPEC strain.

Omptins belonging to the OmpT-like subfamily are known to have subtle differences in substrate specificity (Brannon, Thomassin, et al., 2015; Hwang et al., 2007; McCarter et al., 2004). Studies using peptide libraries to compare OmpP and OmpT activity showed both omptins preferentially cleave substrates between two consecutive basic residues, but that OmpP appears to have a slight preference for Lys in the P and P’ sites (Hwang et al., 2007). In addition to subtle differences in amino acid motif preference, peptide size and secondary structure also impact substrate specificity (Brannon, Thomassin, et al., 2015; Haiko et al., 2009; Hritonenko & Stathopoulos, 2007). For example, AMP α-helicity was shown to be a determining factor for proteolytic activity of the OmpT-like omptin, CroP, from *Citrobacter rodentium* (Brannon, Thomassin, et al., 2015). While ArlC, OmpP and OmpP all readily cleave small unstructured substrates, such as the FRET substrate and C18G, differences in cleavage efficiency were noted for larger or more structured AMPs. OmpP did not cleave Magainin II as efficiently as C18G and did not cleave larger substrates such as mCRAMP, LL-37 and RNase 7 (Figs. 4A, 5AB, 6B). These findings suggest that larger peptides might be excluded from the OmpP active site. While OmpT and ArlC cleaved the FRET substrate, C18G, Magainin II and mCRAMP relatively efficiently, there was a striking difference in LL-37 and RNase 7 cleavage (Figs. 4A, 5A, 6B). Given the similarity in size of mCRAMP and LL-37, and the ability of ArlC to cleave RNase 7, it is unlikely that the 3 amino acid size difference accounts for the marked difference in cleavage efficiency. It is possible that ArlC does not cleave α-helical AMPs, but instead cleaves unstructured and disulfide bond-stabilized peptides. While this possibility requires further study, it is supported by the finding that an *arlC* deletion strain is more susceptible to killing by human defensins (McPhee et al., 2014). Altogether, these findings suggest the presence ArlC and OmpT in the same UPEC isolate may confer a fitness advantage by expanding the spectrum of target substrates.

## 5 Conclusions

Here we show that increased omptin activity is associated with UPEC strains causing symptomatic UTIs. Extensive heterogeneity of omptin activity among UPEC clinical isolates was is due to variations in *ompT* expression and to the presence of a plasmid-encoded *ompT*-like gene *arlC*. Our findings support current profiling practices of UPEC strains that include the *ompT* gene (Johnson & Stell, 2000), but suggest that additional screening for *arlC* should be considered as both genes were exclusively harboured in UPEC strains associated with symptomatic infection. Altogether our findings suggest that the presence of two different omptins in a UPEC strain may provide an additional fitness advantage by expanding the range of AMPs cleaved during UTIs.

## Acknowledgements

This work was supported by the Canadian Institutes of Health Research (CIHR, MOP-15551), the Natural Sciences and Engineering Research Council (NSERC, RGPIN-217482) and the Fonds de Recherche Québec - Nature et Technologies (FRQNT 2013-PR-165926). ID was supported by the Fonds de Recherche Québec - Santé (FRQS). JLT was supported by a Hugh Burke fellowship from the McGill Faculty of Medicine. JLT is supported by a NSERC postdoctoral fellowship and Pasteur-Roux fellowship. JAT and GTM were supported by the Canadian Institutes of Health Research (CIHR, MOP-125998). We thank Dr. Mario Jacques and Mr. Frédéric Berthiaume (Faculté de Médecine Vétérinaire, Université de Montréal) for providing access to the Jasco J-810 spectropolarimeter and technical assistance with CD experiments. We thank Dr. Selena Sagan for the gift of labeling reagents for Southern hybridization. We thank Drs. Olivera Francetic and Yannick Tremblay for helpful comments and suggestions.

## Author contributions

AP, ID, JAT, JLT, JML, and JRB performed experiments. AM, GTM, HLM, ID, JDS, JLT, KD, and SG conceived and designed experiments. HLM, ID, JLT, and SG analyzed the data. HLM and ID wrote early drafts of the manuscript. JLT wrote and reviewed later drafts with support from all other authors.

## Dedication

This publication is dedicated to Dr. Hervé LeMoual who passed away on March 3rd 2018; he was a great mentor that always encouraged his trainees to follow their passions.

## Ethics statement

None required.

