## Appendix for "Identification and characterization of OmpT-like proteases in uropathogenic *Escherichia coli* clinical isolates"

### Appendix: Genome analysis of sequenced Cystitis isolates


| Item | Page |
|---|---|
| Materials and Methods – Genome analysis | A 2 |
| Figure 1. Comparison of cystitis (UTI) isolate genomes with reference strains | A 3 |
| Figure 2. Comparison of plasmid sequences containing pathogenicity island 6 | A 4 |
| Figure 3. Alignment of plasmid 2 from cystitis (UTI) isolate 6 | A 5 |
| Table 1. General features of sequenced cystitis (UTI) isolates | A 6 |
| Table 2. Genome features of sequenced cystitis (UTI) isolates | A 7 |
| Table 3. Predicted genomic islands | A 8 |
| Table 4. Antibiotic resistance genes in cystitis isolates | A 10 |
| Table 4. Unique sequences in cystitis (UTI) isolate 1 | A 18 |
| Table 5. Unique sequences in cystitis (UTI) isolate 6 | A 21 |
| Table 6. Unique sequences in cystitis (UTI) isolate 11 | A 23 |
| References | A 27 |

### Appendix Materials and Methods

#### Genome analysis

Sequenced genomes were annotated using PATRIC (Wattam et al., 2017). Isolate serotypes were determined using the online database SeroTypeFinder (Joensen, Tetzschner, Iguchi, Aarestrup, & Scheutz, 2015). Pathogenicity islands were detected using IslandViewer 4 (Bertelli et al., 2017) and VRprofile 2 (Li et al., 2018). Antibiotic resistance genes were identified in PATRIC, IslandViewer 4, VRprofile 2, BLAST and RGI 4.2.2 CARD 3.0.1 (Jia et al., 2017). Figures of genome alignments and plasmid alignments were generated using BRIGs software (Alikhan, Petty, Ben Zakour, & Beatson, 2011).

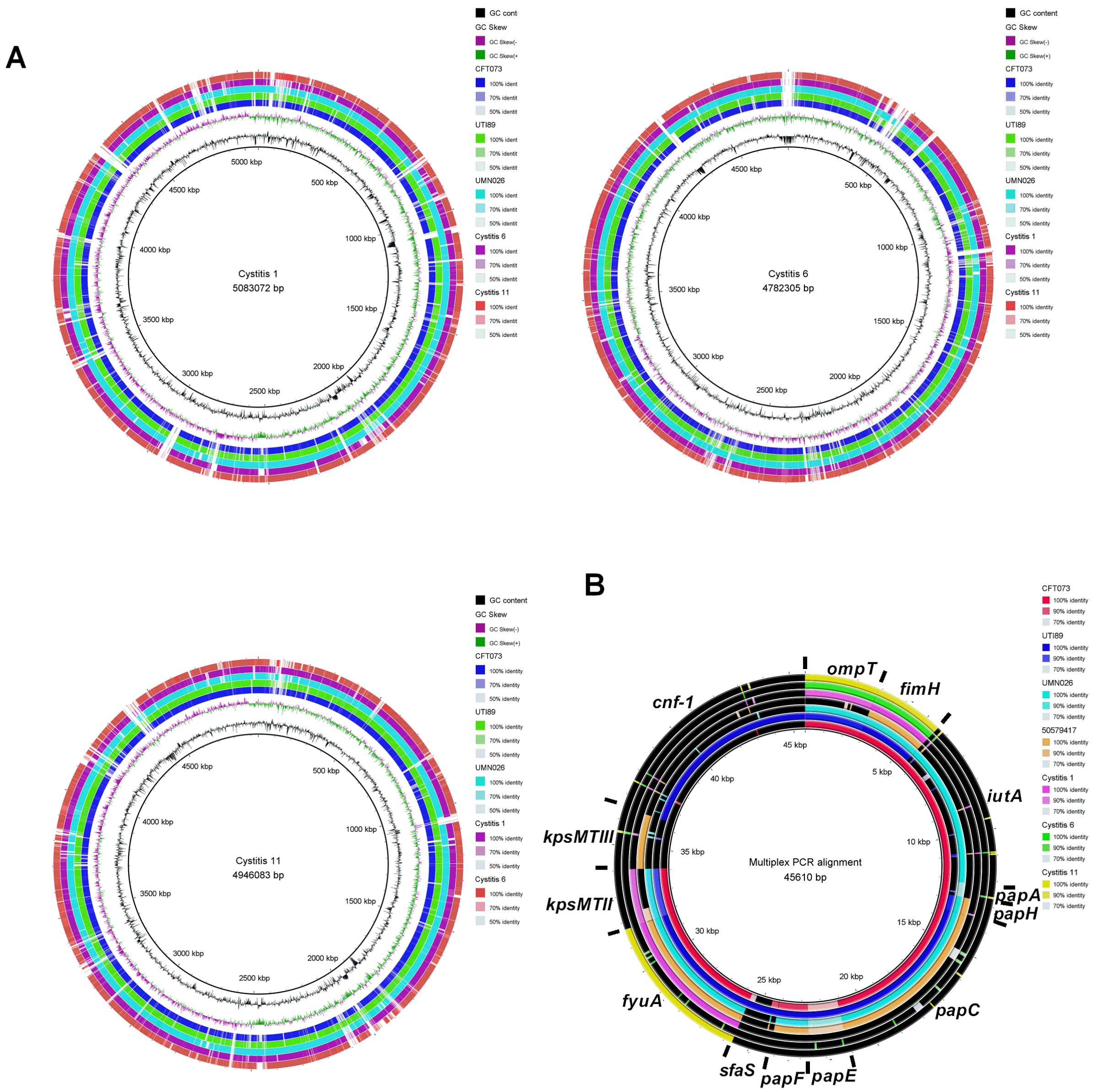


Appendix Figure 1. Comparisons of cystitis (UTI) isolate genomes with reference strains. (A) Indicated cystitis isolates were used as subject sequences in multiple sequence alignments with the indicated UPEC strain genomes using BRIGs software. (B) Genes amplified by multiplex PCR (*ompT, fimH, iutA, papA, papH, papC, papF, fyuA, kpsMTII, papE*, *sfaS*, *kpsMTIII*, *cnf-1*) were used as subject sequences for a multiple sequence alignment of the indicated UPEC strain genome using BRIGs software. Black fill indicates no homology.

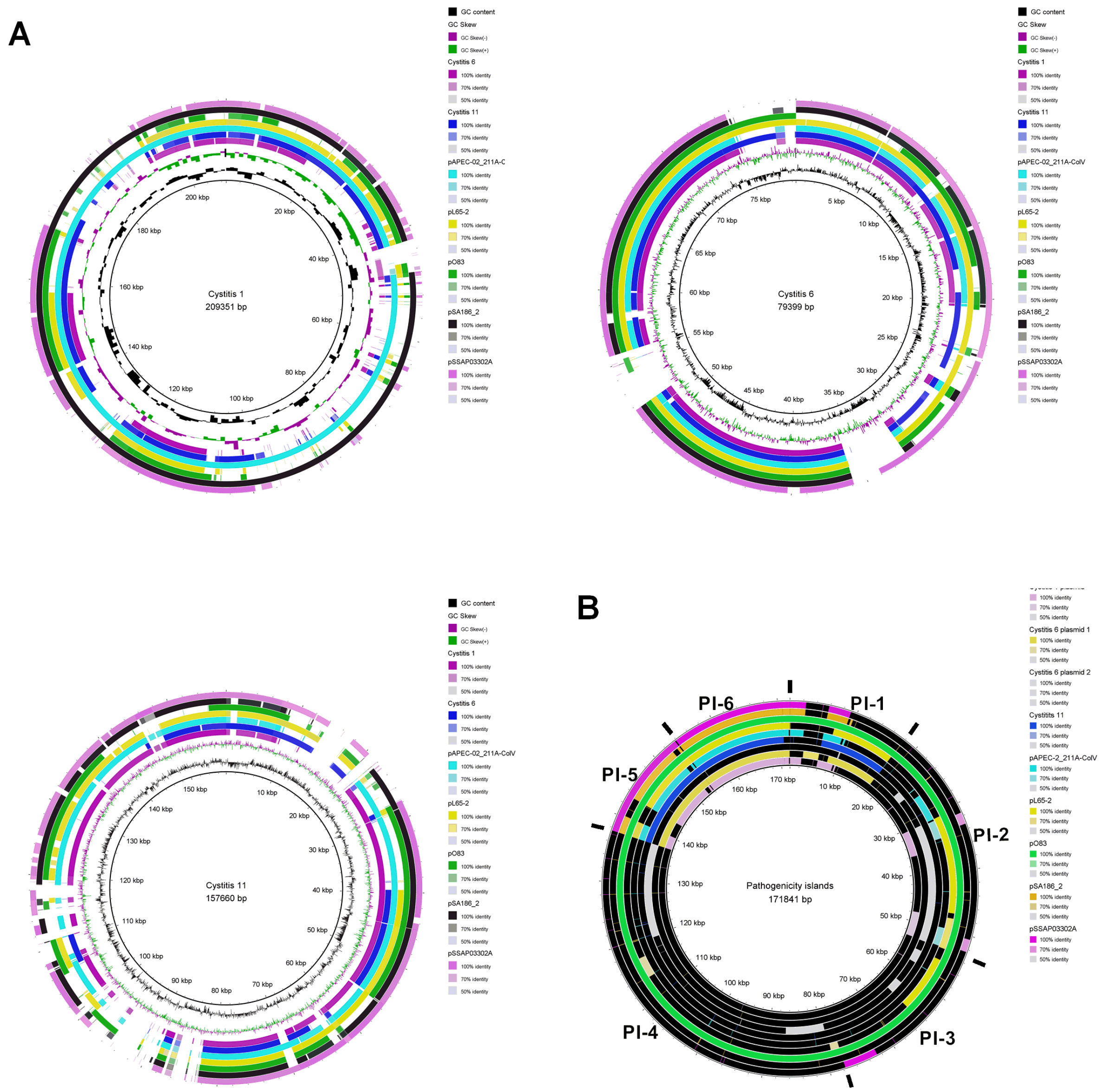


Appendix Figure 2. Comparison of plasmid sequences containing pathogenicity island 6. (A) Plasmids from the indicated cystitis (UTI) isolates were used as subject sequences in multiple sequence alignments with the indicated plasmid containing pathogenicity island 6 from pO83 from *E. coli* NRG857c using BRIGs software. (B) Coding sequences for pathogenicity islands 1, 2, 3, 4, 5, and 6 from pO83 (indicated) were used as subject sequences for a multiple sequence alignment with the indicated plasmid nucleotide sequence using BRIGs software. Black fill indicates no homology.

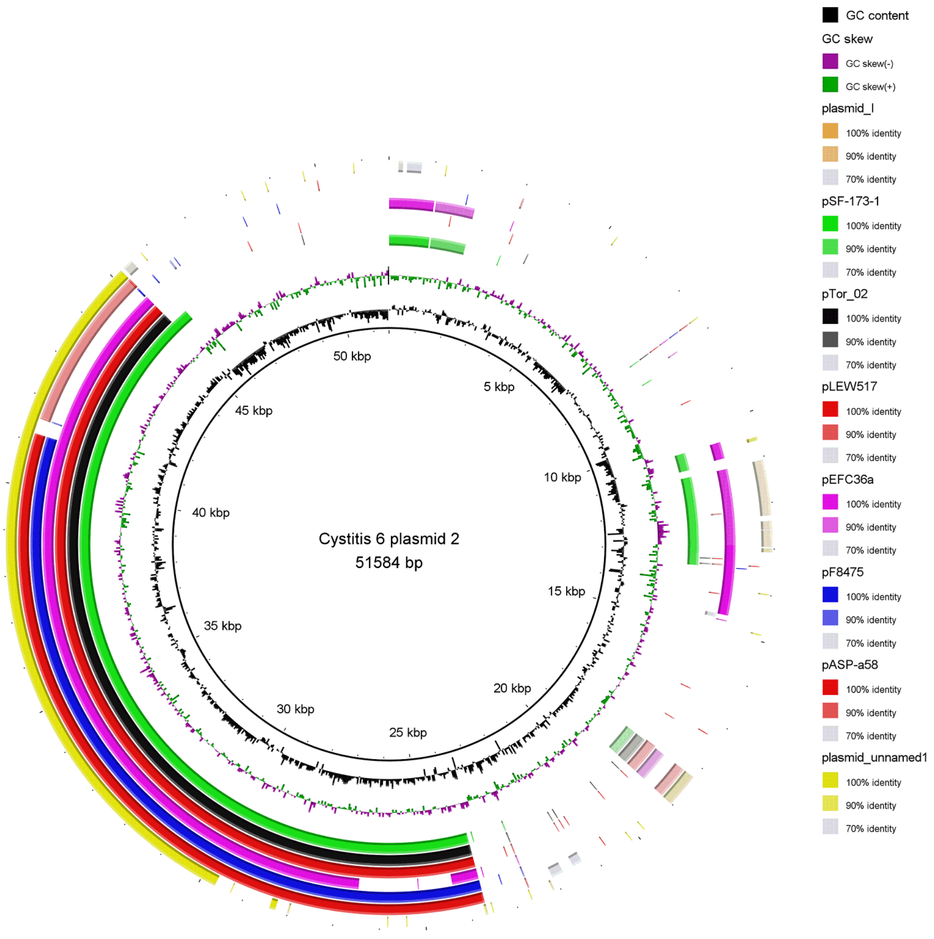


Appendix Figure 3. Alignment of plasmid 2 from cystitis (UTI) isolate 6. Plasmids with sequence homology to plasmid 2 were identified by BLAST search. BRIGs software was used to generate graphical sequence alignments.

Appendix Table 1. General features of sequenced cystitis (UTI) isolates

| Strain | Serotype | Pathotype | Origin/disease | Phylogenetic group | Chromosome | | Plamid | |
|---|---|---|---|---|---|---|---|---|
| | | | | | Size (kb) | G+C content (%) | Size (kb) | G+C content (%) |
| Cystitis 1 | O1:H42 | ExPEC | *Homo sapiens*/ Cystitis | D | 5,107 | 50.5 | 209 | 49.5 |
| Cystitis 6 | O82:H8 | ExPEC | *Homo sapiens*/ Cystitis | B1 | 4,782 | 50.6 | 79, 52 | 48.9, 55.9 |
| Cystitis 11 | O24:H4 | ExPEC | *Homo sapiens*/ Cystitis | D | 4,946 | 50.7 | 157 | 49.8 |

Appendix Table 2. Genome features of sequenced cystitis (UTI) isolates

| Strains | | Cystitis 1 | | Cystitis 6 | | | Cystitis 11 | |
|---|---|---|---|---|---|---|---|---|
| | | Chromosome | Plasmid | Chromosome | Plasmid 1 | Plasmid 2 | Chromosome | Plasmid |
| Number of genes | | 4970 | 301 | 4679 | 137 | 61 | 4875 | 236 |
| rRNA | | 22 | 0 | 22 | 0 | 0 | 22 | 0 |
| tRNA | | 88 | 0 | 90 | 0 | 0 | 89 | 0 |
| Prophages | Complete | 5 | 1 | 0 | 0 | 0 | 5 | 0 |
| | Incomplete | 3 | 0 | 5 | 0 | 0 | 3 | 0 |
| Virulence factors | Number | 173 | 39 | 114 | 14 | 5 | 246 | 26 |
| | % of genes | 3.48 | 12.96 | 2.44 | 10.22 | 8.20 | 5.05 | 11.02 |
| Genomic islands | | 12 | 4 | 11 | 2 | 0 | 15 | 2 |
| Unique sequences[a] | | 73 | N/A | 34 | N/A | N/A | 100 | N/A |

[a]N/A not applicable

Appendix Table 3. Predicted genomic islands

| Isolate | Location | GI[a] | Start | Stop | Size | % GC | # VG[b] | #AMR[c] | tRNA | Features |
|---|---|---|---|---|---|---|---|---|---|---|
| Cystitis 1 | Chromosome | 1 | 108501 | 129938 | 21438 bp | 46.94 | 6 | 0 | seC | GI-like region-1 |
| | | 2 | 147071 | 185155 | 38085 bp | 48.72 | 10 | 0 | 0 | prophage-1 |
| | | 3 | 759876 | 781233 | 21358 bp | 42.64 | 3 | 0 | 0 | GI-like region-2 |
| | | 4 | 1067617 | 1083797 | 16181 bp | 36.70 | 0 | 0 | 0 | T3SS-1 |
| | | 5 | 1974099 | 2012452 | 38354 bp | 48.56 | 14 | 0 | 0 | GI-like region-3 |
| | | 6 | 2496955 | 2544448 | 47494 bp | 48.51 | 0 | 0 | Arg | prophage-2 |
| | | 7 | 2715386 | 2762418 | 47033 bp | 47.59 | 0 | 0 | 0 | prophage-3 |
| | | 8 | 2924030 | 2960151 | 36122 bp | 45.64 | 3 | 0 | 0 | GI-like region-4 |
| | | 9 | 3638710 | 3682745 | 44036 bp | 50.79 | 1 | 1 | 0 | prophage-4 |
| | | 10 | 3972222 | 3989123 | 16902 bp | 56.05 | 0 | 0 | 0 | T6SS-1 |
| | | 11 | 4643625 | 4660145 | 16521 bp | 48.64 | 0 | 0 | 0 | prophage-5 |
| | | 12 | 4802101 | 4835389 | 33289 bp | 51.16 | 1 | 0 | 0 | prophage-6 |
| | Plasmid | 1 | 5292 | 37257 | 31966 bp | 52.16 | 0 | 0 | 0 | T4SS-1 |
| | | 2 | 119770 | 136621 | 16852 bp | 43.93 | 3 | 0 | 0 | prophage-1 |
| | | 3 | 182505 | 194773 | 12269 bp | 49.70 | 2 | 1 | 0 | prophage-2 |
| | | 4 | 200991 | 209349 | 8359 bp | 49.13 | 0 | 0 | 0 | T4SS-2 |
| Cystitis 6 | Chromosome | 1 | 9045 | 26763 | 17719 bp | 40.23 | 0 | 0 | 0 | prophage-1 |
| | | 2 | 457520 | 475359 | 17840 bp | 48.50 | 2 | 0 | 0 | GI-like region-1 |
| | | 3 | 897443 | 919427 | 21985 bp | 45.65 | 0 | 0 | Arg | prophage-2 |
| | | 4 | 1119747 | 1142698 | 22952 bp | 44.95 | 0 | 0 | Thr | prophage-3 |
| | | 5 | 1951044 | 1969169 | 18126 bp | 46.63 | 1 | 0 | 0 | prophage-4 |
| | | 6 | 2249278 | 2256258 | 6981 bp | 44.33 | 2 | 0 | 0 | prophage-5 |
| | | 7 | 2316877 | 2325663 | 8787 bp | 52.88 | 0 | 0 | 0 | GI-like region-2 |
| | | 8 | 2695320 | 2714439 | 19120 bp | 50.54 | 9 | 0 | 0 | GI-like region-3 |
| | | 9 | 2728674 | 2734357 | 5684 bp | 52.48 | 1 | 0 | Leu | prophage-6 |
| | | 10 | 4256804 | 4265449 | 8646 bp | 34.65 | 0 | 0 | 0 | T3SS-1 |
| | | 11 | 4460470 | 4468593 | 8124 bp | 48.25 | 0 | 0 | 0- | prophage-7 |
| | Plasmid 1 | 1 | 3 | 12486 | 12484 bp | 51.11 | 0 | 0 | 0 | T4SS-1 |
| | | 2 | 15576 | 46399 | 30824 bp | 49.91 | 9 | 0 | 0 | prophage-1 |
| | Plasmid 2 | 1 | 29675 | 30495 | 821 bp | 55.9 | 0 | 0 | 0 | IS15 |

| | | | | | | | | | | |
|---|---|---|---|---|---|---|---|---|---|---|
| | | 2 | 33726 | 34605 | 880 bp | 60.5 | 0 | 0 | 0 | IS6100 |
| | | 3 | 42222 | 45221 | 3000 bp | 63.2 | 0 | 0 | 0 | ISPa38 |
| Cystitis 11 | Chromosome | 1 | 109066 | 132168 | 23103 bp | 47.29 | 7 | 0 | seC | GI-like region-1 |
| | | 2 | 1044432 | 1070691 | 26260 bp | 54.86 | 0 | 0 | 0 | T6SS-1 |
| | | 3 | 1856379 | 1874039 | 17661 bp | 36.50 | 6 | 1 | 0 | GI-like region-2 |
| | | 4 | 1894582 | 1918137 | 23556 bp | 51.16 | 6 | 0 | 0 | GI-like region-3 |
| | | 5 | 1970393 | 2014554 | 44162 bp | 51.03 | 0 | 0 | Gly | prophage-1 |
| | | 6 | 2069555 | 2104371 | 34817 bp | 51.43 | 0 | 0 | 0 | prophage-2 |
| | | 7 | 2436915 | 2489158 | 52244 bp | 48.83 | 0 | 0 | 0 | prophage-3 |
| | | 8 | 2835003 | 2844947 | 9945 bp | 48.52 | 1 | 0 | 0 | prophage-4 |
| | | 9 | 3120766 | 3148412 | 27647 bp | 51.06 | 1 | 2 | 0 | prophage-5 |
| | | 10 | 3246346 | 3305235 | 58890 bp | 49.39 | 2 | 0 | 0 | prophage-6 |
| | | 11 | 3371214 | 3382861 | 11648 bp | 48.20 | 0 | 0 | 0 | GI-like region-4 |
| | | 12 | 3825971 | 3850884 | 24914 bp | 49.38 | 8 | 0 | 0 | GI-like region-5 |
| | | 13 | 4115749 | 4151643 | 35895 bp | 50.78 | 3 | 1 | 0 | GI-like region-6 |
| | | 14 | 4119559 | 4168240 | 48682 bp | 50.17 | 1 | 0 | Leu | prophage-7 |
| | | 15 | 4403545 | 444486 | 40942 bp | 43.08 | 7 | 0 | 0 | GI-like region-7 |
| | Plasmid | 1 | 5280 | 23125 | 17846 bp | 54.73 | 2 | 1 | 0 | prophage-1 |
| | | 2 | 125191 | 157658 | 32468 bp | 53.35 | 0 | 0 | 0 | T4SS-1 |

[a]Genomic island
[b]VG indicates known virulence genes
[c]Antimicrobial resistance genes

Appendix Table 4. Antibiotic resistance genes in cystitis (UTI) isolates

| Isolate | Location | Gene | Function; Resistance mechanism | Resistance to |
|---|---|---|---|---|
| Cystitis 1 | Chromosome | *acrA* | Subunit of an RND efflux pump; antibiotic efflux | Aminoglycosides |
| | | *acrD* | Part of an RND efflux pump; antibiotic efflux | Aminoglycosides |
| | | *acrE* | Part of AcrEF-TolC efflux pump; antibiotic efflux | Fluoroquinolones, cephamycin, cephalosporin, penam |
| | | *acrF* | Part of AcrEF-TolC efflux pump; antibiotic efflux | Fluoroquinolones, cephamycin, cephalosporin, penam |
| | | *adeF* | Membrane fusion protein of the multidrug efflux complex AdeFGH; antibiotic efflux | Fluoroquinolone, tetracycline |
| | | *ampC* | Enzymatic degradation of ß-lactam rings; antibiotic inactivation | Broad and extended spectrum ß-lactamases |
| | | *cmeB* | Inner membrane transporter in CmeABC RND efflux channel; antibiotic efflux | Cephalosporins, macrolides, fluoroquinolones, fusidic acid |
| | | *cmeC* | Outer membrane channel in CmeABC RND efflux channel; antibiotic efflux | Cephalosporins, macrolides, fluoroquinolones, fusidic acid |
| | | *cyaA* | Adenylate cyclase variant S352T; antibiotic target alteration | Fosfomycin |
| | | *emrA* | Membrane fusion protein in EmrAB-TolC efflux pump complex; antibiotic efflux | Fluoroquinolone |
| | | *emrB* | Translocase in EmrAB-TolC efflux pump complex; antibiotic efflux | Fluoroquinolone |
| | | *emrD* | Multidrug transporter that couples efflux of amphipathic compounds with proton import across the plasma membrane; antibiotic efflux | Detergents |
| | | *emrE* | Small multidrug resistance efflux; antibiotic efflux | Macrolides |
| | | *emrY* | Multidrug transport across the inner membrane; antibiotic efflux | Tetracycline |
| | | *ermK* | Membrane fusion protein that works with ErmY and TolC as part of a MFS efflux pump; antibiotic efflux | Tetracycline |
| | | *ftsI* | Sequence variant D350N, S357N of PBP3; antibiotic target alteration | Cephamycin, cephalosporin, penam, carbapenam, monobactam |
| | | *glpT* | Sequence variant E448K of the active importer GlpT; antibiotic target alteration | Fosfomycin |
| | | *macA* | Membrane fusion protein that acts with MacB and TolC to form an ABC antibiotic efflux comples; antibiotic efflux | Macrolides |
| | | *macB* | ABC transporter that acts with MacA and TolC to form an | Macrolides (14-/15-membered lactones) |

| | | |
|---|---|---|
| | ABC antibiotic efflux comples; antibiotic efflux | |
| *marA* | Regulates MDR efflux pump and regulates porin synthesis; reduced antibiotic permeability, antibiotic efflux | Tetracycline, penem, penam, carbapenem, cephamycin, cephalosporin, rifamycin, phenicol, monobactam, glycycline, fluoroquinolone, triclosan |
| *marR* | MarR variant G103S Y137H causes efflux pump overexpression; antibiotic target alteration, antibiotic efflux | Tetracylcines, penam, cephalosporins, glycycline, rifamycin, phenicol, triclosan, fluoroquinolones |
| *mdfA* | Multidrug efflux pump; antibiotic efflux | Tetracycline, benzalkonium chloride, rhodamine |
| *mdtA* | Membrane fusion protein RND efflux pump; antibiotic efflux | Aminocoumarins |
| *mdtB* | Transporter that forms multimeric complex with MdtC; antibiotic efflux | Aminocoumarins |
| *mdtC* | Transporter that forms multimeric complex with MdtB; antibiotic efflux | Aminocoumarins |
| *mdtD* | MFS transporter; antibiotic efflux | Aminocoumarins |
| *mdtE* | Membrane fusion protein that works with MdtF and TolC as part of a MFS efflux pump; antibiotic efflux | Penam, fluoroquinolones, macrolides |
| *mdtF* | Inner membrane transporter that works with MdtE and TolC as part of a MFS efflux pump; antibiotic efflux | Penam, fluoroquinolones, macrolides |
| *mdtH* | MFS transporter; antibitioc efflux | fluoroquinolones |
| *mdtM* | MFS transporter; antibiotic efflux | Nucleosides, phenicol, lincosamides, fluoroquinolones, acridine dye |
| *mdtN* | Part of MdtNOP MFS efflux pump; antibiotic efflux | Nucleoside antibitiocs, acridine dye |
| *mdtO* | Part of MdtNOP MFS efflux pump; antibiotic efflux | Nucleoside antibitiocs, acridine dye |
| *mdtP* | Part of MdtNOP MFS efflux pump; antibiotic efflux | Nucleoside antibitiocs, acridine dye |
| *msbA* | Multidrug resistance transporter homologue; antibiotic efflux | Nitroimidazole |
| *nfsA* | Variant Y45C of major oxygen insensitive nitroreductase in *Escherichia coli;* antibiotic target alteration | Nitrofuran |
| *pmrD* | Histidine kinase involved in regulation of polymyxin resistance; antibiotic target alteration | Polymyxins and peptide antibiotics |
| *pmrF* | Glycosyl transferase; antibiotic target alteration | Polymyxins and peptide antibiotics |
| *pmrH* | UDP-4-amino-4-deoxy-L-arabinose-oxoglutarate aminotransferase; antibiotic target alteration | Polymyxins and peptide antibiotics |
| *pmrI* | UDP-4-amino-4deoxy-L-arabinose formyltransferase; antibiotic target alteration | Polymyxins and peptide antibiotics |

| | | | | |
|---|---|---|---|---|
| | | *pmrJ* | Catalyzes deformylation of L-Ara4-formyL-*N*; antibiotic target alteration | Polymyxins and peptide antibiotics |
| | | *pmrK* | Undecaprenyl phosphate-alpha-4-amino-4-deoxy-L-arabinosyl transferase; antibiotic target alteration | Polymyxins and peptide antibiotics |
| | | *pmrL* | Sucrose-6 phosphate hydrolase; antibiotic target alteration | Polymyxins and peptide antibiotics |
| | | *pmrM* | Subunit of undecaprenyl phosphate-alpha-L-Ara4N flippase; antibiotic target alteration | Polymyxins and peptide antibiotics |
| | | *sapA* | Periplasmic solute binding protein; antibiotic efflux | Antimicrobial peptides |
| | | *sapB* | Permease subunit; antibiotic efflux | Antimicrobial peptides |
| | | *sapC* | Permease subunit; antibiotic efflux | Antimicrobial peptides |
| | | *sapD* | ATPase; antibiotic efflux | Antimicrobial peptides |
| | | *sapF* | ATPase; antibiotic efflux | Antimicrobial peptides |
| | | *tufA* | Sequence variant R234F of elongation factor Tu; antibiotic target alteration | Pulvomycin, elfamycin |
| Cystitis 6 | Chromosome | *acrA* | subunit of AcrAB-TolC RND efflux pump; antibiotic efflux | Tetracycline, penam, cephalosporin, rifamycin, phenicol, glycycline, fluoroquinolone, triclosan |
| | | *acrB* | subunit of AcrAB-TolC RND efflux pump; antibiotic efflux | Tetracycline, penam, cephalosporin, rifamycin, phenicol, glycycline, fluoroquinolone, triclosan |
| | | *acrD* | Part of an RND efflux pump; antibiotic efflux | Aminoglycosides |
| | | *acrE* | Part of AcrEF-TolC RND efflux pump; antibiotic efflux | Cephamycin, cpehalosporin, penam, fluoroquinolone |
| | | *acrF* | Part of AcrEF-TolC RND efflux pump; antibiotic efflux | Cephamycin, cpehalosporin, penam, fluoroquinolone |
| | | *adeF* | Membrane fusion protein of the AdeFGH RND efflux pump; antibiotic efflux | Tetracycline, fluoroquinolones |
| | | *ampC* | Enzymatic degradation of ß-lactam rings; antibiotic inactivation | Broad and extended spectrum ß-lactamases |
| | | *bcr* | Part of an efflux system; antibiotic efflux | Bicyclomycins |
| | | *cmeB* | Inner membrane transporter of the CmeABC RND efflux complex; antibiotic efflux | Macrolides, cephalosporins, fusidic acid, fluoroquinolones |
| | | *emrA* | Part of the EmrAB-TolC MFS efflux pump; antibiotic efflux | Fluoroquinolones |
| | | *emrB* | Part of the EmrAB-TolC MFS efflux pump; antibiotic efflux | Fluoroquinolones |
| | | *emrK* | Part of the EmKY-TolC MFS efflux pump; antibiotic efflux | Tetracyclines |
| | | *emrY* | Part of the EmKY-TolC MFS efflux pump; antibiotic efflux | Tetracyclines |
| | | *ermK* | Erm 23S rRNA methyltransferase; antibiotic target | Lincosamides, macrolides, streptogramins |

| | | |
|---|---|---|
| | alteration | |
| *ftsI* | Sequence variant D350N, S357N of PBP3; antibiotic target alteration | Cephamycin, cephalosporin, penam, carbapenam, monobactam |
| *gyrA* | Point mutation (S83L); antibiotic target modification | Fluoroquinolones, nybomycin |
| *marA* | Regulates MDR efflux pump and regulates porin synthesis; reduced antibiotic permeability, antibiotic efflux | Tetracycline, penem, penam, carbapenem, cephamycin, cephalosporin, rifamycin, phenicol, monobactam, glycycline, fluoroquinolone, triclosan |
| *marR* | Regulates *marAB* operon; antibiotic target alteration, antibiotic efflux | Tetracylcines, penam, cephalosporins, glycycline, rifamycin, phenicol, triclosan, fluoroquinolones |
| *marR* | MarR variant G103S Y137H causes efflux pump overexpression; antibiotic target alteration, antibiotic efflux | Tetracylcines, penam, cephalosporins, glycycline, rifamycin, phenicol, triclosan, fluoroquinolones |
| *mdfA* | Multidrug efflux pump; antibiotic efflux | Tetracycline, benzalkonium chloride, rhodamine |
| *mdtA* | Membrane fusion protein RND efflux pump; antibiotic efflux | Aminocoumarins |
| *mdtB* | Transporter that forms multimeric complex with MdtC; antibiotic efflux | Aminocoumarins |
| *mdtC* | Transporter that forms multimeric complex with MdtB; antibiotic efflux | Aminocoumarins |
| *mdtE* | Membrane fusion protein that works with MdtF and TolC as part of a MFS efflux pump; antibiotic efflux | Penam, fluoroquinolones, macrolides |
| *mdtF* | Inner membrane transporter that works with MdtE and TolC as part of a MFS efflux pump; antibiotic efflux | Penam, fluoroquinolones, macrolides |
| *mdtH* | MFS transporter; antibitioc efflux | fluoroquinolones |
| *mdtK* | Part of a multidrug and toxic compounds extrusions transporter; antibiotic efflux | Norfloxacin, doxorubicin, acriflavine |
| *mdtM* | MFS transporter; antibiotic efflux | Nucleosides, phenicol, lincosamides, fluoroquinolones, acridine dye |
| *mdtN* | Part of MdtNOP MFS efflux pump; antibiotic efflux | Nucleoside antibitiocs, acridine dye |
| *mdtO* | Part of MdtNOP MFS efflux pump; antibiotic efflux | Nucleoside antibitiocs, acridine dye |
| *mdtP* | Part of MdtNOP MFS efflux pump; antibiotic efflux | Nucleoside antibitiocs, acridine dye |
| *msbA* | Multidrug resistance transporter homologue; antibiotic efflux | Nitroimidazole |
| *nfsA* | Variant Y45C of major oxygen insensitive nitroreductase in *Escherichia coli;* antibiotic target alteration | Nitrofuran |
| *pmrD* | Histidine kinase involved in regulation of polymyxin | Polymyxins and peptide antibiotics |

| | | | | |
|---|---|---|---|---|
| | | | resistance; antibiotic target alteration | |
| | | *pmrF* | Glycosyl transferase; antibiotic target alteration | Polymyxins and peptide antibiotics |
| | | *pmrH* | UDP-4-amino-4-deoxy-L-arabinose-oxoglutarate aminotransferase; antibiotic target alteration | Polymyxins and peptide antibiotics |
| | | *pmrI* | UDP-4-amino-4deoxy-L-arabinose formyltransferase; antibiotic target alteration | Polymyxins and peptide antibiotics |
| | | *pmrJ* | Catalyzes deformylation of L-Ara4-formyL-*N*; antibiotic target alteration | Polymyxins and peptide antibiotics |
| | | *pmrK* | Undecaprenyl phosphate-alpha-4-amino-4-deoxy-L-arabinosyl transferase; antibiotic target alteration | Polymyxins and peptide antibiotics |
| | | *pmrL* | Sucrose-6 phosphate hydrolase; antibiotic target alteration | Polymyxins and peptide antibiotics |
| | | *pmrM* | Subunit of undecaprenyl phosphate-alpha-L-Ara4N flippase; antibiotic target alteration | Polymyxins and peptide antibiotics |
| | | *tufA* | Sequence variant R234F of elongation factor Tu; antibiotic target alteration | Pulvomycin, elfamycin |
| | Plasmid 1 | *cmeC* | Outer membrane channel of the CmeABC RND antibiotic efflux pump; antibiotic efflux | Cephalosporin, macrolides, fluoroquinolones, fusidic acid |
| | | *macA* | Membrane fusion protein that acts with MacB and TolC to form an ABC antibiotic efflux comples; antibiotic efflux | Macrolides |
| | | *macB* | ABC transporter that acts with MacA and TolC to form an ABC antibiotic efflux comples; antibiotic efflux | Macrolides (14-/15-membered lactones) |
| | | *mdtH* | MFS antibtiotic efflux pump; antibiotic efflux | Fluoroquinolones |
| | Plasmid 2 | *aadA* | Aminoglycoside nucleotidyltransferase, antibiotic inactivation | Aminoglycosides |
| | | *dfrA12* | Dihydrofolate reductase; antibiotic target replacement | Diaminopyrimidine |
| | | *mphA* | Macrolide 2’-phosphotransferase; antibiotic inactivation | Macrolides; preferentially inactivates 14-membered macrolides over 16-membered macrolides |
| | | *sul1* | Dihydropteroate synthase type-2; antibiotic target replacement | Sulfonamides, sulfone |
| | | *tetA* | Tetracycline efflux protein TetA; antibiotic efflux | Tetracyclines, Glycylcycline |
| Cystitis 11 | Chromosome | *acrA* | subunit of AcrAB-TolC RND efflux pump; antibiotic efflux | Tetracycline, penam, cephalosporin, rifamycin, phenicol, glycycline, fluoroquinolone, triclosan |
| | | *acrB* | subunit of AcrAB-TolC RND efflux pump; antibiotic efflux | Tetracycline, penam, cephalosporin, rifamycin, phenicol, glycycline, fluoroquinolone, triclosan |
| | | *acrD* | RND antibiotic efflux pump; antibiotic efflux | Aminoglycosides |

| | | |
|---|---|---|
| *acrE* | Membrane fusion protein of a RND efflux transporter; antibiotic efflux | Penam, cephamycin, cephalosporin, fluoroquinolones |
| *acrF* | Inner membrane transporter component of a RND efflux transporter; antibiotic efflux | Penam, cephamycin, cephalosporin, fluoroquinolones |
| *adeF* | Membrane fusion protein of the AdeFGH RND efflux pump; antibiotic efflux | Tetracycline, fluoroquinolones |
| *ampC* | Class C ß-lactamase; antibiotic inactivation | Broad and extended spectrum cephalosporins |
| *cmeB* | Inner membrane transporter in CmeABC RND efflux channel; antibiotic efflux | Cephalosporins, macrolides, fluoroquinolones, fusidic acid |
| *cmeC* | Outer membrane channel in CmeABC RND efflux channel; antibiotic efflux | Cephalosporins, macrolides, fluoroquinolones, fusidic acid |
| *cyaA* | Adenylate cyclase variant S352T; antibiotic target alteration | Fosfomycin |
| *emrA* | Part of the EmrAB-TolC MFS efflux pump; antibiotic efflux | Fluoroquinolones |
| *emrB* | Part of the EmrAB-TolC MFS efflux pump; antibiotic efflux | Fluoroquinolones |
| *emrD* | Multidrug transporter that couples efflux of amphipathic compounds with proton import across the plasma membrane; antibiotic efflux | Detergents |
| *emrE* | Small MDR transporter; antibiotic efflux | Macrolides |
| *emrK* | Part of the EmKY-TolC MFS efflux pump; antibiotic efflux | Tetracyclines |
| *emrY* | Part of the EmKY-TolC MFS efflux pump; antibiotic efflux | Tetracyclines |
| *emrY* | MFS antibiotic efflux pump; antibiotic efflux | Tetracyclines |
| *ermA* | RNA methylase; antibiotic target alteration | Macrolides, streptogramins, lincosamides |
| *ermB* | RNA methylase; antibiotic target alteration | Macrolides, streptogramins, lincosamides |
| *ermK* | RNA methylase; antibiotic target alteration | Macrolides, streptogramins, lincosamides |
| *ftsI* | Sequence variant D350N, S357N of PBP3; antibiotic target alteration | Cephamycin, cephalosporin, penam, carbapenam, monobactam |
| *glpT* | Sequence variant E448K of the active importer GlpT; antibiotic target alteration | Fosfomycin |
| *macA* | Membrane fusion protein that acts with MacB and TolC to form an ABC antibiotic efflux comples; antibiotic efflux | Macrolides |
| *macB* | ABC transporter that acts with MacA and TolC to form an ABC antibiotic efflux comples; antibiotic efflux | Macrolides (14-/15-membered lactones) |
| *marA* | Global activator protein that induces MDR efflux and | Tetracycline, penem, penam, carbapenem, cephalosporin, |

| | | |
|---|---|---|
| | downregulates OmpF synthesis; reduced permeability to antibiotic, antibiotic efflux | rifamycin, phenicol, monobactam, glycycline, fluoroquinolone, triclosan |
| *marR* | MarR variant G103S Y137H causes efflux pump overexpression; antibiotic target alteration, antibiotic efflux | Tetracylcines, penam, cephalosporins, glycycline, rifamycin, phenicol, triclosan, fluoroquinolones |
| *mdfA* | Multidrug efflux pump; antibiotic efflux | Tetracycline, benzalkonium chloride, rhodamine |
| *mdtA* | Membrane fusion component of the MdtABC RND efflux pump; antibiotic efflux | Aminocoumarin resistance |
| *mdtB* | Transporter in the MdtABC RND efflux pump; antibiotic efflux | Aminocoumarin resistance |
| *mdtC* | Transporter in the MdtABC RND efflux pump; antibiotic efflux | Aminocoumarin resistance |
| *mdtE* | Membrane fusion protein of a RND efflux transporter; antibiotic efflux | Penam, macrolides, fluoroquinolones |
| *mdtF* | Multidrug inner membrane transporter of an RND efflux transporter; antibiotic efflux | Penam, macrolides, fluoroquinolones |
| *mdtH* | MFS transporter; antibitioc efflux | fluoroquinolones |
| *mdtM* | MFS transporter; antibiotic efflux | Nucleosides, phenicol, lincosamides, fluoroquinolones, acridine dye |
| *mdtN* | Predicted inner membrane fusion protein of MFS efflux pump; antibiotic efflux | Nucleoside antibiotics, acridine dye |
| *mdtO* | Uncharacterized component of MFS efflux pump; antibiotic efflux | Nucleoside antibiotics, acridine dye |
| *mdtP* | Predicted outer membrane component of MFS efflux pump; antibiotic efflux | Nucleoside antibiotics, acridine dye |
| *msbA* | Multidrug resistance transporter homologue; antibiotic efflux | Nitroimidazole |
| *nfsA* | Variant Y45C of major oxygen insensitive nitroreductase in *Escherichia coli;* antibiotic target alteration | Nitrofuran |
| *pmrD* | Histidine kinase involved in regulation of polymyxin resistance; antibiotic target alteration | Polymyxins and peptide antibiotics |
| *pmrF* | Glycosyl transferase; antibiotic target alteration | Polymyxins and peptide antibiotics |
| *pmrH* | UDP-4-amino-4-deoxy-L-arabinose-oxoglutarate aminotransferase; antibiotic target alteration | Polymyxins and peptide antibiotics |
| *pmrI* | UDP-4-amino-4deoxy-L-arabinose formyltransferase; antibiotic target alteration | Polymyxins and peptide antibiotics |

| | | | |
|---|---|---|---|
| | *pmrJ* | Catalyzes deformylation of L-Ara4-formyL-*N*; antibiotic target alteration | Polymyxins and peptide antibiotics |
| | *pmrK* | Undecaprenyl phosphate-alpha-4-amino-4-deoxy-L-arabinosyl transferase; antibiotic target alteration | Polymyxins and peptide antibiotics |
| | *pmrL* | Sucrose-6 phosphate hydrolase; antibiotic target alteration | Polymyxins and peptide antibiotics |
| | *pmrM* | Subunit of undecaprenyl phosphate-alpha-L-Ara4N flippase; antibiotic target alteration | Polymyxins and peptide antibiotics |
| | *sapA* | Periplasmic solute binding protein; antibiotic efflux | Antimicrobial peptides |
| | *sapB* | Permease subunit; antibiotic efflux | Antimicrobial peptides |
| | *sapC* | Permease subunit; antibiotic efflux | Antimicrobial peptides |
| | *sapD* | ATPase; antibiotic efflux | Antimicrobial peptides |
| | *sapF* | ATPase; antibiotic efflux | Antimicrobial peptides |
| | *tetA* | Tetracycline efflux protein TetA; antibiotic efflux | Tetracyclines, Glycylcycline |
| | *tetB* | Part of MFS efflux pump; antibiotic efflux | Tetracycline, doxycycline, minocycline |
| | *tetC* | Part of MFS efflux pump; antibiotic efflux | Tetracycline |
| | *tufA* | Sequence variant R234F of elongation factor Tu; antibiotic target alteration | Pulvomycin, elfamycin |
| Plasmid | *cmeC* | Outer membrane channel of the CmeABC RND antibiotic efflux pump; antibiotic efflux | Cephalosporin, macrolides, fluoroquinolones, fusidic acid |
| | *macA* | Membrane fusion protein that acts with MacB and TolC to form an ABC antibiotic efflux comples; antibiotic efflux | Macrolides |
| | *macB* | ABC transporter that acts with MacA and TolC to form an ABC antibiotic efflux comples; antibiotic efflux | Macrolides (14-/15-membered lactones) |

Appendix Table 5. Unique sequences in cystitis (UTI) isolate 1

| Start | End | Length[a] | In GI[b] | Number of genes | Description[c] |
|---|---|---|---|---|---|
| 92527 | 99018 | 6492 | Y | 8 | Hypothetical protein, Hypothetical protein, Transcriptional regulator LacI family, PTS system IIA component, putative sugar phosphoesterase component IIB, Putative integral membrane protein, Transketolase N-terminal section, Transketolase C-terminal section |
| 99269 | 109488 | 10220 | Y/N/Y | | Hypothetical protein, Hypothetical protein, Cobalt-zinc-cadmium resistance protein CzcA/ Cation efflux system protein CusA, Hypothetical protein, Hypothetical protein, Periplasmic lysozyme inhibitor of c-type lysozyme, Hypothetical protein, Hypothetical protein, N-acetylmannosamine-6-phosphate 2-epimerase, PTS system/ maltose and glucose-specific IIC component, RpiR family transcriptional regulator, putative exported protein, probable transposase, |
| 111022 | 111909 | 888 | Y | 1 | Hypothetical protein |
| 119743 | 125490 | 5748 | Y | 3 | Mobile element protein, Mobile element protein, AidA-I adhesin-like protein |
| 171411 | 173689 | 2279 | Y | 2 | Beta-1,4-galactosyltransferase, O-antigen ligase |
| 205046 | 206569 | 1524 | Y | 2 | Hypothetical protein, Hypothetical protein |
| 207862 | 215371 | 7510 | Y/N | 7 | DsORF-h1, Hypothetical protein, core protein, Hypothetical protein, Hypothetical protein, core protein, Hypothetical protein |
| 223771 | 225181 | 1411 | N | 2 | Prophage Lp2 protein 6, Hypothetical protein |
| 757555 | 764928 | 7374 | Y | 7 | Integrase, Hypothetical protein, tRNA-dihydrouridine synthase, Hypothetical protein, Putative DNA binding protein, hypothetical protein, HigA (antitoxin to HigB) |
| 767309 | 768408 | 1100 | Y | 2 | Hypothetical protein, Hypothetical protein |
| 769304 | 770308 | 1005 | Y | 1 | Transposase |
| 771382 | 773282 | 1901 | Y | 2 | Hypothetical protein, Hypothetical protein |
| 773582 | 781971 | 8390 | Y | 6 | Type I restriction-modification system/ DNA-methyltransferase subunit M, Anticodon nuclease, Type I restriction-modification system/ specificity subunit S, hypothetical protein, Type I restriction-modification system/ restriction subunit R, hypothetical protein |
| 861933 | 863466 | 1534 | N | 2 | hypothetical protein, hypothetical protein |
| 929045 | 937462 | 8418 | N | 6 | hypothetical protein, hypothetical protein, hypothetical protein, Mobile element protein, Beta-1,3-glucosyltransferase, UDP-glucose 6-dehydrogenase |
| 1197765 | 1206221 | 8457 | Y | 8 | CRISPR-associated helicase Cas3, hypothetical protein, CRISPR-associated protein/Cse1 family, CRISPR-associated protein/Cse2 family, CRISPR-associated protein/ Cse4 family, CRISPR-associated protein/ Cas5e family, CRISPR-associated protein/Cse3 family, CRISPR-associated protein Cas1 |
| 1499352 | 1501563 | 2212 | N | 2 | hypothetical protein, LPS glycosyltransferase |
| 1606008 | 1607967 | 1960 | N | 4 | Minor fimbrial subunit StfE, Minor fimbrial subunit StfF, Minor fimbrial subunit StfG, Uncharacterized protein YadU in stf fimbrial cluster |
| 1868446 | 1869415 | 970 | N | 1 | Uncharacterized protein YehA precursor |
| 1945389 | 1952171 | 6783 | N | 2 | Membrane protein involved in the export of O-antigen, UDP-N-acetylglucosamine 2-epimerase |
| 2002072 | 2007197 | 5126 | Y | 3 | Aspartate ammonia-lyase, Tripeptide aminopeptidase, Anaerobic C4-dicarboxylate transporter DcuC |
| 2008196 | 2010951 | 2756 | Y | 4 | hypothetical protein, Isoaspartyl aminopeptidase, hypothetical protein, Transposase |
| 2013119 | 2017875 | 4757 | Y | 3 | Mg(2+)-transport-ATPase-associated protein MgtC, Inosine-uridine preferring nucleoside hydrolase, Transporter/MFS |

| | | | | | |
|---|---|---|---|---|---|
| | | | | | superfamily |
| 2027105 | 2034581 | 7477 | Y | 5 | Putative transcriptional regulator LYSR-type, Aspartate racemase, Anaerobic C4-dicarboxylate transporter, Aspartate ammonia-lyase, Anaerobic C4-dicarboxylate transporter DcuB |
| 2035649 | 2044005 | 8357 | Y | 4 | hypothetical protein, Fumarate respiration transcriptional regulator DcuR, regulatory protein GntR, Anaerobic C4-dicarboxylate transporter |
| 2135213 | 2136774 | 1562 | N | 1 | Flagellar hook-associated protein FliD |
| 2137384 | 2137941 | 558 | N | 1 | Flagellar biosynthesis protein FliC |
| 2148532 | 2149345 | 814 | N | 1 | hypothetical protein |
| 2150523 | 2163017 | 12495 | Y | 6 | Type I restriction-modification system/DNA-methyltransferase subunit M, Type I restriction-modification system/ specificity subunit S, hypothetical protein, hypothetical protein, Type I restriction-modification system/ restriction subunit R, Putative predicted metal-dependent hydrolase |
| 2187720 | 2190160 | 2441 | N | 3 | hypothetical protein, Deoxyguanosinetriphosphate triphosphohydrolase, dNTP triphosphohydrolase (putative) |
| 2502136 | 2502681 | 546 | Y | 1 | hypothetical protein |
| 2505785 | 2507465 | 1681 | Y | 1 | hypothetical protein |
| 2512412 | 2513766 | 1355 | Y | 2 | TolA protein, hypothetical protein |
| 2538335 | 2538862 | 528 | Y | 0 | part of a phage tail fiber protein |
| 2640841 | 2641887 | 1047 | Y | 1 | hypothetical protein |
| 2643180 | 2646480 | 3301 | N | 3 | VgrG protein, hypothetical protein, hypothetical protein |
| 2697571 | 2698622 | 1052 | N | 3 | Phenylacetic acid degradation protein PaaY, Phenylacetic acid degradation operon negative regulatory protein PaaX, Transcriptional activator feaR |
| 2715468 | 2716403 | 936 | Y | 1 | Phage tail fiber protein |
| 2724014 | 2741946 | 17933 | Y/N/Y | 20 | Phage tail length tape-measure protein 1, Phage tail length tape-measure protein 1, hypothetical protein, hypothetical protein, hypothetical protein, hypothetical protein, **Conserved hypothetical protein**, hypothetical protein, hypothetical protein, hypothetical protein, hypothetical protein, hypothetical protein, NAD+--asparagine ADP-ribosyltransferase, 62kDa structural protein, Putative phage terminase, hypothetical protein, hypothetical protein, hypothetical protein, Chromosome (plasmid) partitioning protein ParB, Trk system potassium uptake protein TrkG |
| 2743886 | 2745867 | 1982 | Y | 2 | hypothetical protein, hypothetical protein |
| 2751431 | 2754932 | 3502 | Y | 5 | Bacteriophage-encoded homolog of DNA replication protein DnaC, hypothetical protein, hypothetical protein, hypothetical protein, Rac prophage repressor |
| 2755245 | 2756003 | 759 | Y | 1 | Superinfection exclusion protein B |
| 2756171 | 2757337 | 1167 | Y | 3 | Kil protein, Putative bacteriophage protein, Phage protein |
| 2757792 | 2758520 | 729 | N | 1 | Exodeoxyribonuclease VIII |
| 2758798 | 2762186 | 3389 | N | 6 | Exodeoxyribonuclease VIII, Recombinational DNA repair protein RecT, hypothetical protein, Phage protein, ydaQ protein, Putative lambdoid prophage Rac integrase |
| 2920575 | 2940019 | 19445 | Y | 15 | Integrase, hypothetical protein, hypothetical protein, hypothetical protein, hypothetical protein, hypothetical protein, unknown (no homologous in databases), DNA helicase, hypothetical protein, hypothetical protein, Transposase, hypothetical protein, hypothetical protein, hypothetical protein, hypothetical protein, |
| 2941285 | 2960445 | 19161 | Y | 14 | hypothetical protein, hypothetical protein, Site-specific recombinase XerD, DNA repair protein RadC, hypothetical protein, hypothetical protein, Transcriptional regulator/ (Cro/CI family), hypothetical protein, hypothetical protein, hypothetical |

| | | | | | |
|---|---|---|---|---|---|
| | | | | | protein, hypothetical protein, hypothetical protein, type III restriction enzyme (res subunit), hypothetical protein |
| 3618000 | 3627112 | 9113 | N | 5 | VgrG protein, hypothetical protein, Rhs family protein, hypothetical protein, Rhs family protein |
| 3670522 | 3672667 | 2146 | Y | 2 | hypothetical protein, hypothetical protein |
| 3676650 | 3677582 | 933 | Y | 1 | hypothetical protein |
| 3850940 | 3852767 | 1828 | N | 0 | |
| 3972235 | 3979319 | 7085 | N | 5 | Protein ImpG/VasA, Uncharacterized protein ImpH/VasB, Type VI secretion lipoprotein/VasD, Uncharacterized protein ImpJ/VasE, Outer membrane protein ImpK/VasF (OmpA/MotB domain) |
| 3982676 | 3986259 | 3584 | Y | 1 | IcmF-related protein |
| 3987795 | 3990829 | 3035 | Y | 2 | Secreted protein Hcp, hypothetical protein |
| 4043652 | 4044493 | 842 | N | 1 | Phosphoesterase |
| 4059435 | 4060801 | 1367 | N | 1 | Ferric hydroxamate outer membrane receptor FhuA |
| 4072250 | 4078414 | 6165 | N | 6 | Chaperone protein EcpD, Outer membrane usher protein HtrE, Fimbrial protein YadM, Fimbrial protein YadL, Fimbrial protein YadK, Fimbrial protein YadC |
| 4291685 | 4294387 | 2703 | N | 2 | hypothetical protein, Type I restriction-modification system/ restriction subunit R |
| 4303111 | 4305804 | 2694 | N | 2 | hypothetical protein, hypothetical protein |
| 4307776 | 4313434 | 5659 | N | 1 | adherence and invasion outermembrane protein (Inv, enhances Peyer's patches colonization) |
| 4338354 | 4370454 | 32101 | N/Y | 11 | transposase and inactivated derivative, hypothetical protein, hypothetical protein, putative DNA helicase, putative RNA helicase, Type II restriction enzyme/methylase subunits, putative ATP-dependent helicase, hypothetical protein, hypothetical protein, putative membrane protein, Outer membrane protein and related peptidoglycan-associated (lipo)proteins, hypothetical protein |
| 4371197 | 4371767 | 571 | Y | 1 | ORF25 |
| 4373966 | 4383447 | 9482 | Y/N | 7 | hypothetical protein, hypothetical protein, hypothetical protein, hypothetical protein, hypothetical protein, hypothetical protein, hypothetical protein |
| 4405084 | 4405646 | 563 | N | 1 | hypothetical protein |
| 4773665 | 4781054 | 7390 | N | 5 | IS/phage/ Transposon-related functions, hypothetical protein, hypothetical protein, core protein, core protein, |
| 4808730 | 4810063 | 1334 | N | 2 | hypothetical protein, hypothetical protein |
| 4810567 | 4811397 | 831 | N | 1 | Phage tail fibers |
| 4815114 | 4816052 | 939 | Y | 1 | hypothetical protein |
| 4825453 | 4828858 | 3406 | Y | 3 | hypothetical protein, hypothetical protein, DNA-cytosine methyltransferase |
| 4833607 | 4835504 | 1898 | N | 2 | Cox, C protein |
| 4874755 | 4875788 | 1034 | N | 1 | Inner membrane protein |
| 4881491 | 4887240 | 5750 | N | 6 | Putative cytoplasmic protein, hypothetical protein, Putative membrane-associated metal-dependent hydrolase, Hypothetical radical SAM family enzyme, hypothetical protein, Putative hydrolase |
| 4952534 | 4953592 | 1059 | N | 1 | Transposase |
| 4977905 | 4978540 | 636 | N | 1 | hypothetical protein |

[a]bp
[b]Y: yes, N: no; Blue corresponds to the blue gene description
[c] Red indicates half the coding sequence is present; Bold indicates internal deletion; blue corresponds to the blue GI location

Appendix Table 6. Unique sequences in cystitis (UTI) isolate 6

| Start | End | Length[a] | In GI[b] | Number of genes | Description[c] |
|---|---|---|---|---|---|
| 1 | 17937 | 17937 | Y | 17 | hypothetical protein, hypothetical protein, Putative cytoplasmic protein USSDB7A, Arylsulfatase regulator, Radical SAM domain heme biosynthesis protein, Radical SAM domain heme biosynthesis protein, His-Xaa-Ser repeat protein, hypothetical protein, hypothetical protein, Chromosome (plasmid) partitioning protein ParB, Recombinase, hypothetical protein, hypothetical protein, hypothetical protein, hypothetical protein, hypothetical protein, Integrase |
| 19533 | 23218 | 3686 | Y | 3 | Phage DNA transfer protein, Phage DNA transfer protein, regulatory protein |
| 23528 | 25533 | 2006 | Y | 1 | Phage tail fibers |
| 362972 | 371780 | 8809 | Y | 9 | Asparagine synthetase [glutamine-hydrolyzing], Asparagine synthetase [glutamine-hydrolyzing], Glycerol-3-phosphate cytidylyltransferase, Membrane protein involved in the export of O- antigen, hypothetical protein, hypothetical protein, hypothetical protein, hypothetical protein, Putative N-acetylgalactosaminyl-diphosphoundecaprenol glucuronosyltransferase |
| 434684 | 447623 | 12940 | N/Y | 11 | hypothetical protein, hypothetical protein, Predicted ATP-dependent endonuclease of the OLD family, ATP-dependent DNA helicase UvrD/PcrA, hypothetical protein, hypothetical protein, hypothetical protein, Exonuclease SbcC, hypothetical protein, hypothetical protein, hypothetical protein |
| 461321 | 473333 | 12013 | N/Y | 14 | hypothetical protein, hypothetical protein, probable monooxygenase, Cysteinyl-tRNA synthetase, Zinc uptake regulation protein ZUR, Putative metal chaperone/ involved in Zn homeostasis/ GTPase of COG0523 family, C4-type zinc finger protein(DksA/TraR family), GTP cyclohydrolase I, GTP cyclohydrolase I, Carbonic anhydrase (gamma class), NADPH dependent preQ0 reductase, putative inner membrane protein, Manganese ABC transporter/ inner membrane permease protein SitD, Manganese ABC transporter/inner membrane permease protein SitC |
| 475741 | 491876 | 16136 | N/Y/N | 16 | Manganese ABC transporter/ATP-binding protein SitB, Manganese ABC transporter/periplasmic-binding protein SitA, Threonyl-tRNA synthetase, peptidase, Dihydroorotase, Porphobilinogen synthase, hypothetical protein, FAD dependent oxidoreductase, hypothetical protein, putative secreted protein, Zinc ABC transporter/ periplasmic-binding protein ZnuA, Zinc ABC transporter/ inner membrane permease protein ZnuB, Zinc ABC transporter/ inner membrane permease protein ZnuB, ABC transporter/ATP-binding protein, Putative phosphatase, Putative phosphatase, |
| 535212 | 538414 | 3203 | N | 2 | Flagellar hook-associated protein FliD, Flagellar biosynthesis protein FliC |
| 908378 | 909732 | 1355 | Y | 2 | TolA protein, hypothetical protein |
| 911128 | 912495 | 1368 | Y | 1 | probable bacteriophage protein STY2043 |
| 914322 | 915218 | 897 | Y | 1 | hypothetical protein |
| 967813 | 968868 | 1056 | Y | 1 | Mobile element protein |
| 1117605 | 1119765 | 2161 | Y | 1 | Bicyclomycin resistance protein |
| 1121023 | 1123899 | 2877 | Y | 2 | Chromosome (plasmid) partitioning protein ParB, Trk system potassium uptake protein TrkG |
| 1125106 | 1125829 | 724 | Y | 0 | |
| 1125990 | 1129600 | 3611 | Y | 5 | Phage antitermination protein, IS/ phage/Transposon-related functions, hypothetical bacteriophage protein, Putative cytoplasmic protein, hypothetical protein |
| 1130306 | 1131315 | 1010 | Y | 2 | hypothetical protein, LygF |
| 1131764 | 1135077 | 3314 | N | 4 | Bacteriophage-encoded homolog of DNA replication protein DnaC, hypothetical protein, hypothetical protein, Regulatory |

| | | | | | |
|---|---|---|---|---|---|
| | | | | | protein Cro of bacteriophage BP-933W |
| 1135233 | 1136281 | 1049 | N | 2 | Phage protein, IS/ phage/Transposon-related functions |
| 1136449 | 1137615 | 1167 | N | 2 | Kil protein, Phage protein, |
| 1138070 | 1138816 | 747 | N | 1 | Exodeoxyribonuclease VIII |
| 1139077 | 1142466 | 3390 | N | 5 | Exodeoxyribonuclease VIII, Recombinational DNA repair protein RecT, Phage protein, ydaQ protein, Putative lambdoid prophage Rac integrase |
| 1256434 | 1257373 | 940 | N | 4 | transposase and inactivated derivative, hypothetical protein, Transposase, Transposase |
| 1439363 | 1441005 | 1643 | N | 2 | Uncharacterized protein YcdU, Uncharacterized protein YmdE |
| 2252604 | 2256411 | 3808 | N | 3 | hypothetical protein, hypothetical protein, Phage integrase/ Phage P4-associated |
| 2631438 | 2632843 | 1406 | N | 2 | hypothetical protein, hypothetical protein/ Unknown protein (Putative secreted protein) |
| 2638026 | 2638530 | 505 | N | 0 | |
| 2707045 | 2709006 | 1962 | Y | 3 | hypothetical protein, hypothetical protein, unknown |
| 2729916 | 2731171 | 1256 | N | 1 | Integrase |
| 2802563 | 2805725 | 3163 | N | 3 | TRAP-type transport system/small permease component/ predicted N-acetylneuraminate transporter, TRAP-type C4-dicarboxylate transport system/ large permease component, TRAP-type C4-dicarboxylate transport system/ periplasmic component |
| 4241941 | 4254806 | 12866 | Y | 17 | hypothetical protein, hypothetical protein, resolvase, hypothetical protein, hypothetical protein, hypothetical protein, hypothetical protein, diguanylate cyclase/phosphodiesterase (GGDEF & EAL domains) with PAS/PAC sensor(s), hypothetical protein, hypothetical protein, hypothetical protein, hypothetical protein, hypothetical protein, hypothetical protein, hypothetical protein, hypothetical protein, Integrase |
| 4757988 | 4778127 | 20140 | Y | 27 | hypothetical protein, hypothetical protein, DNA-binding protein H-NS, hypothetical protein, hypothetical protein, hypothetical protein, hypothetical protein, GALNS arylsulfatase regulator (Fe-S oxidoreductase), hypothetical protein, hypothetical protein, thiJ/pfpI family protein, hypothetical protein, NUDIX hydrolase, Transcriptional regulator (AraC family), Integral membrane protein, hypothetical protein, Transcriptional regulator (AlpA like), hypothetical protein, hypothetical protein, hypothetical protein, Integrase/recombinase (XerC/CodV family), putative enzyme; Integration, recombination (Phage or Prophage Related), putative enzyme; Integration, recombination (Phage or Prophage Related), putative enzyme; Integration, recombination (Phage or Prophage Related), hypothetical protein, hypothetical protein, Transposase and inactivated derivatives |
| 4781408 | 4782305 | 898 | Y | 1 | hypothetical protein |

[a] bp

[b] Y: yes, N: no; Blue corresponds to the blue gene description

[c] Blue corresponds to the blue GI location

Appendix Table 7. Unique sequences in cystitis (UTI) isolate 11

| Start | End | Length[a] | In GI[b] | Number of genes | Description[c] |
|---|---|---|---|---|---|
| 53318 | 54471 | 1154 | Y | 1 | Hypothetical protein |
| 60486 | 61814 | 1329 | Y | 1 | Mobile element protein |
| 66554 | 71961 | 5408 | Y | 6 | Hypothetical protein, Hypothetical protein, DNA binding protein H-NS homolog, Hypothetical protein, dNTP triphosphohydrolase, Hemolysin E |
| 93640 | 100131 | 6492 | N/Y | 8 | Hypothetical protein, Hypothetical protein, Transcriptional regulator LacI family, PTS system IIA component, putative sugar phosphoesterase component IIB, Putative integral membrane protein, Transketolase N-terminal section, Transketolase C-terminal section |
| 100382 | 109054 | 8673 | Y/N | 10 | Cobalt-zinc-cadmium resistance protein CzcA/ Cation efflux system protein CusA, Hypothetical protein, Hypothetical protein, Periplasmic lysozyme inhibitor of c-type lysozyme, Hypothetical protein, Hypothetical protein, N-acetylmannosamine-6-phosphate 2-epimerase, PTS system maltose and glucose-specific IIC component, RpiR family transcriptional regulator, putative exported protein |
| 110386 | 111721 | 1336 | Y | 4 | Mobile element protein, probable transposase, Transposase, Hypothetical protein |
| 113255 | 114142 | 888 | Y | 1 | Hypothetical protein |
| 121972 | 127726 | 5755 | Y | 3 | Mobile element protein, Mobile element protein, AidA-I adhesin-like protein |
| 171434 | 173712 | 2279 | Y | 2 | Beta-1,4-galactosyltransferase, O-antigen ligase |
| 261819 | 263138 | 1320 | N | 2 | Putative cytoplasmic protein, Mobile element protein |
| 380154 | 380822 | 669 | N | 2 | Prevent host death protein/Phd antitoxin, Death on curing protein/ Doc toxin |
| 436499 | 437810 | 1312 | N | 2 | Mobile element protein, Putative cytoplasmic protein |
| 510751 | 512074 | 1324 | N | 1 | Mobile element protein |
| 522438 | 523766 | 1329 | Y | 1 | Mobile element protein |
| 789229 | 797600 | 8372 | Y | 8 | Hypothetical protein, Glycerol kinase, Hypothetical protein, Ribokinase, Hypothetical protein, ADP-ribosylglycohydrolase, Fatty acyl responsive regulator, Possible GPH family transporter for arabinosides |
| 841386 | 842713 | 1328 | N | 1 | Mobile element protein |
| 880627 | 881954 | 1328 | N | 1 | Mobile element protein |
| 1049588 | 1050827 | 1240 | N | 1 | Hypothetical protein |
| 1057872 | 1059240 | 1369 | N | 2 | Hypothetical protein, Hypothetical protein |
| 1234072 | 1235400 | 1329 | N | 1 | Mobile element protein |
| 1457321 | 1458649 | 1329 | N | 1 | Putative outer membrane protein |
| 1524259 | 1525075 | 817 | N | 1 | Uncharacterized protein YadU in stf fimbrial cluster |
| 1764883 | 1766325 | 1443 | N | 1 | Molybdate metabolism regulator |
| 1773822 | 1775150 | 1329 | Y | 1 | Mobile element protein |
| 1779718 | 1780687 | 970 | Y | 1 | Uncharacterized protein YehA precursor |
| 1854559 | 1868386 | 13828 | Y | 9 | putative glycosyl transferase, Mobile element protein, Mobile element protein, Sialic acid biosynthesis protein NeuD/ O-acetyltransferase, N-acetylneuraminate synthase, N-Acetylneuraminate cytidylyltransferase, UDP-N-acetylglucosamine 2-epimerase, hypothetical protein, N-acetylneuraminic acid synthase-like protein |
| 1869519 | 1871201 | 1683 | Y | 1 | Glycosyl transferase group 1 |

| | | | | | |
|---|---|---|---|---|---|
| 1978623 | 1979489 | 867 | N | 1 | putative transcriptional regulator |
| 1979791 | 1980456 | 666 | N | 1 | Phage or Prophage Related |
| 1980905 | 1981663 | 759 | N | 1 | Bacteriophage-encoded homolog of DNA replication protein DnaC |
| 1984213 | 1984864 | 652 | N | 0 | |
| 1989870 | 1990900 | 1031 | Y | 2 | Hypothetical protein, Hypothetical protein |
| 2012030 | 2013277 | 1248 | Y | 1 | Phage tail fiber protein |
| 2049480 | 2050358 | 879 | Y | 1 | Mobile element protein |
| 2055676 | 2057239 | 1564 | N | 1 | Flagellar hook-associated protein FliD |
| 2068610 | 2069434 | 825 | N | 0 | |
| 2070171 | 2071982 | 1812 | N | 1 | Hypothetical protein |
| 2078841 | 2079758 | 918 | N | 1 | Serine acetyltransferase |
| 2094001 | 2095196 | 1196 | N | 1 | Hypothetical protein |
| 2099069 | 2100098 | 1030 | N | 2 | Hypothetical protein, Hypothetical protein |
| 2101327 | 2104477 | 3151 | N | 6 | putative DNA-binding protein, hypothetical protein, Transcriptional regulator/XRE family, putative membrane protein, unknown protein encoded by prophage CP-933T, putative integrase |
| 2129647 | 2130975 | 1329 | N | 2 | Hypothetical protein, Flagellar biosynthesis protein FlhB |
| 2248120 | 2249448 | 1329 | N | 1 | Mobile element protein |
| 2254799 | 2256127 | 1329 | N | 1 | Mobile element protein |
| 2265562 | 2266882 | 1321 | N | 2 | Excinuclease cho (excinuclease ABC alternative C subunit), Mobile element protein |
| 2442097 | 2442642 | 546 | Y | 2 | Hypothetical protein, IS/phage/ Transposon-related functions |
| 2445725 | 2447404 | 1680 | Y | 1 | Hypothetical protein |
| 2521789 | 2523116 | 1328 | Y | 1 | Mobile element protein |
| 2536073 | 2537396 | 1324 | N | 1 | Mobile element protein |
| 2574125 | 2577574 | 3450 | N | 2 | Putative hydrolase, Hypothetical protein |
| 2578274 | 2588113 | 9840 | N | 8 | no significant similarities, hypothetical protein, hypothetical protein, Rhs-family protein, VgrG protein, hypothetical protein, hypothetical protein, hypothetical protein |
| 2633948 | 2635276 | 1329 | Y | 1 | Mobile element protein |
| 2671799 | 2673127 | 1329 | N | 2 | Oxidoreductase (putative), Mobile element protein |
| 2744476 | 2745596 | 1121 | N | 2 | Hypothetical protein, Hypothetical protein |
| 2854519 | 2855846 | 1328 | Y | 1 | Mobile element protein |
| 2897568 | 2898896 | 1329 | N | 1 | Mobile element protein |
| 2962930 | 2964240 | 1311 | N | 2 | Putative cytoplasmic protein, Mobile element protein |
| 3119888 | 3120871 | 984 | N | 0 | |
| 3121457 | 3122590 | 1134 | N | 1 | Prophage Clp protease-like protein |
| 3122678 | 3123968 | 1291 | N | 2 | Hypothetical protein, Hypothetical protein |
| 3124678 | 3127448 | 2771 | N | 3 | Zinc binding domain / DNA primase /Phage P4-associated / Replicative helicase RepA/ Pha, Hypothetical protein, Hypothetical protein |
| 3194818 | 3195987 | 1170 | N | 2 | Putative ATPase component of ABC transporter with duplicated ATPase domain, L,D-transpeptidase YbiS |
| 3250851 | 3252098 | 1248 | N | 1 | Phage tail fiber protein |

| | | | | | |
|---|---|---|---|---|---|
| 3268304 | 3270333 | 2030 | Y | 2 | Hypothetical protein, Hypothetical protein |
| 3270590 | 3272684 | 2095 | Y | 3 | Predicted transcriptional regulator, Hypothetical protein, Phage major capsid protein |
| 3274952 | 3279544 | 4593 | N | 9 | Hypothetical protein, Hypothetical protein, Hypothetical protein, Hypothetical protein, IS/ phage/ Transposon-related functions, Hypothetical protein, Hypothetical protein, Phage DNA binding protein, site-specific recombinase/phage integrase family |
| 3294991 | 3295695 | 705 | Y | 2 | Regulatory protein cro, Phage repressor |
| 3296108 | 3298426 | 2319 | Y | 3 | Hypothetical protein, Phage antitermination protein N, Hypothetical protein |
| 3302188 | 3303631 | 1444 | Y | 2 | Eae protein, Hypothetical protein |
| 3312404 | 3313731 | 1328 | N | 1 | Mobile element protein |
| 3371556 | 3373208 | 1653 | Y | 1 | Hypothetical protein |
| 3374508 | 3375901 | 1394 | Y | 3 | no significant similarities, Hypothetical protein, Rhs-family protein |
| 3377200 | 3383693 | 6494 | Y | 3 | Transposase, Transposase, Hypothetical protein |
| 3391993 | 3394128 | 2136 | N | 3 | Hypothetical protein, Ornithine decarboxylase, Mobile element protein |
| 3397651 | 3398212 | 562 | N | 2 | Hypothetical protein, ABC-type sugar transport system/periplasmic component |
| 3607995 | 3609323 | 1329 | N | 1 | Mobile element protein |
| 3757998 | 3758611 | 614 | Y | 1 | Hypothetical protein |
| 3784880 | 3786207 | 1328 | Y | 1 | Mobile element protein |
| 3827613 | 3830272 | 2660 | Y | 2 | Mobile element protein, Mobile element protein |
| 3903105 | 3903954 | 850 | N | 1 | Phosphoesterase |
| 3918822 | 3920188 | 1367 | N | 1 | Ferric hydroxamate outer membrane receptor FhuA |
| 4139527 | 4141169 | 1643 | Y | 1 | Hypothetical protein |
| 4143342 | 4145291 | 1950 | Y | 1 | Hypothetical protein |
| 4146944 | 4147626 | 683 | N | 1 | Phage-related protein |
| 4156103 | 4157607 | 1505 | N | 1 | Hypothetical protein |
| 4176846 | 4182742 | 5897 | N | 5 | Transcriptional regulator/RpiR family, Pantothenate:Na+ symporter, Bona fide RidA/YjgF/TdcF/RutC subgroup, D-aminoacylase, D-serine deaminase |
| 4186367 | 4194686 | 8320 | N | 5 | Hypothetical protein, Hypothetical protein, Hypothetical protein, Type I restriction-modification system/ specificity subunit S, Type I restriction-modification system/ specificity subunit R |
| 4229745 | 4230315 | 571 | Y | 1 | ORF25 |
| 4258051 | 4259379 | 1329 | N | 1 | Mobile element protein |
| 4288044 | 4289271 | 1228 | N | 1 | Hypothetical protein |
| 4295450 | 4296851 | 1402 | N | 1 | Hypothetical protein |
| 4382926 | 4399461 | 16536 | Y/N | 11 | conserved protein of unknown function, Hypothetical protein, Hypothetical protein, Hypothetical protein, DNA sulfur modification protein DndE, DNA sulfur modification protein DndD, 3'-phosphoadenosine 5'-phosphosulfate sulfurtransferase DndC, DNA sulfur modification protein DndB, Hypothetical protein, Hypothetical protein, Hypothetical protein |
| 4401865 | 4411010 | 9146 | Y | 12 | Mobile element protein, LysR family transcriptional regulator YeiE, Hypothetical protein, Sodium/glutamate symport protein, Hypothetical protein, Hypothetical protein, Hypothetical protein, Transcriptional regulator/ ArsR family, Transcriptional regulator/ TetR family, Tetracycline efflux protein TetA, Hypothetical protein, Right origin-binding protein, Mobile element protein |

| | | | | | |
|---|---|---|---|---|---|
| 4411949 | 4413800 | 1852 | Y | 2 | Transposase, Hypothetical protein |
| 4415069 | 4417517 | 2449 | Y | 3 | Hypothetical protein, Hypothetical protein, Hypothetical protein |
| 4418723 | 4423934 | 5212 | Y | 5 | Hypothetical protein, Hypothetical protein, Hypothetical protein, conserved hypothetical protein, Hypothetical protein |
| 4426949 | 4428102 | 1154 | Y | 1 | Hypothetical protein |
| 4428676 | 4432267 | 3592 | N | 3 | Mobile element protein, Hypothetical protein, Hypothetical protein |
| 4496164 | 4497492 | 1329 | N | 1 | Mobile element protein |
| 4831272 | 4832424 | 1153 | N | 1 | Possible exported protein |

[a]bp
[b]Y: yes, N: no; Blue corresponds to the blue gene description
[c] Blue corresponds to the blue GI location
